# HAF Prevents Hepatocyte Apoptosis and Hepatocellular Carcinoma through Transcriptional Regulation of the NF-κB pathway

**DOI:** 10.1101/2024.01.09.574894

**Authors:** Karen Acuña Pilarte, Ethan Conrad Reichert, Yangsook Song Green, Lily Marie-Therese Halberg, Sydney A. McFarland, Patrice N. Mimche, Martin Golkowski, Severin Donald Kamdem, Kathleen M. Maguire, Scott. A. Summers, J. Alan Maschek, Jordan William Reelitz, James Eric Cox, Kimberley Jane Evason, Mei Yee Koh

## Abstract

**Background:** Hepatocellular carcinoma (HCC) incidence is increasing worldwide due to the obesity epidemic, which drives metabolic dysfunction-associated steatohepatitis (MASH) that can lead to HCC. However, the molecular pathways that lead to MASH-HCC are poorly understood. We have previously reported that male mice with global haploinsufficiency of hypoxia-associated factor, HAF (*SART1*^*+/*-^) spontaneously develop MASH/HCC. However, the cell type(s) responsible for HCC associated with HAF loss are unclear.

**Results:** *SART1*-floxed mice were crossed with mice expressing Cre-recombinase within hepatocytes (Alb-Cre; hepS^-/-^) or macrophages (LysM-Cre, macS^-/-^). Only hepS^-/-^ mice (both male and female) developed HCC suggesting that HAF protects against HCC primarily within hepatocytes. HAF-deficient macrophages showed decreased P-p65 and P-p50 and in many major components of the NF-κB pathway, which was recapitulated using HAF siRNA *in vitro*. HAF depletion increased apoptosis both *in vitro* and *in vivo*, suggesting that HAF mediates a tumor suppressor role by suppressing hepatocyte apoptosis. We show that HAF regulates NF-κB activity by controlling transcription of *TRADD* and *RIPK1*. Mice fed a high-fat diet (HFD) showed marked suppression of HAF, P-p65 and TRADD within their livers after 26 weeks, but manifest profound upregulation of HAF, P-65 and TRADD within their livers after 40 weeks of HFD, implicating deregulation of the HAF-NF-κB axis in the progression to MASH. In humans, HAF was significantly decreased in livers with simple steatosis but significantly increased in HCC compared to normal liver.

**Conclusions:** HAF is novel transcriptional regulator of the NF-κB pathway that protects against hepatocyte apoptosis and is a key determinant of cell fate during progression to MASH and MASH-HCC.

## Introduction

Hepatocellular carcinoma (HCC) is the most common type of liver malignancy and represents the fourth leading cause of cancer-related deaths worldwide (1-3). HCC typically arises in the context of liver damage resulting from excessive inflammation associated with risk factors such as metabolic dysfunction-associated steatotic liver disease (MASLD), infection with hepatitis B or C virus (HBV and HCV respectively) and excessive alcohol consumption (1). Although HBV infection is the most common cause of HCC, MASLD has become the fastest growing etiology for HCC in the United States due to the increasing prevalence of obesity (4-7). MASLD is defined as an excess of intrahepatic lipid storage that can progress to steatohepatitis (MASH), a condition associated with steatosis and chronic inflammation that promotes cell death and regeneration resulting in hepatic injury (5, 8), (9-11). MASH may advance to fibrosis and cirrhosis, the most prominent cause of HCC (9). However, unlike that typically observed in patients with HBC/HCV infection, tumorigenesis in MASH patients may also occur without progression to cirrhosis and the molecular mechanisms by which MASH contributes to hepatocarcinogenesis remains poorly understood (12).

MASH-related chronic hepatic inflammation is associated with excessive cell death (12, 13). The tumor necrosis factor alpha (TNF) is a central cytokine that regulates cell survival and cell death in hepatocytes. TNF stimulates the activation of the nuclear factor kB (NF-κB) pathway by binding to the TNFR1 receptor and subsequently triggering the recruitment of TRADD (TNFRSF1A Associated Via Death Domain), RIPK1 (Receptor Interacting Serine/Threonine Kinase 1), TRAF2 (TNF Receptor Associated Factor 2) and cIAPs (Cellular Inhibitor of Apoptosis Proteins) to the signaling complex I (14, 15). Assembly of complex I mediates the recruitment of other proteins such as TAK1 (Transforming growth factor-β activated kinase 1) and NEMO (NF-kB Essential Modifier) to facilitate proteasomal degradation of IκBα (NF-κB Inhibitor Alpha), an inhibitor of the p50/p65 heterodimer, transcriptional mediators of the NF-κB pathway. Upon release, p50/p65 are translocated into the nucleus promoting transcriptional activation of cell survival, proliferation, differentiation and pro-inflammatory genes, as well as apoptosis regulators (16). TNF stimulation of the TNFR1 receptor can also trigger apoptosis and/or necroptosis by recruiting caspase-8 and RIPK3, respectively, in cells with impaired NF-κB activation (17). Thus, NF-κB plays an important homeostatic role in hepatocytes by activating both anti-apoptotic and pro-inflammatory responses to cellular stresses, thus ensuring that hepatocytes are protected from cell death while appropriate inflammatory responses are initiated (18).

The hypoxia-associated factor, HAF (encoded by the gene *SART1*) is a multifunctional molecule that was initially identified as a transcriptional co-activator of *EPO* (19). We have shown that HAF similarly binds and activates HIF-2α dependent transcription of *VEGFA* and *OCT3/4* (19, 20). In addition to its DNA binding co-transcriptional activity, HAF also functions as an E3 ubiquitin ligase that ubiquitinates and degrades HIF-1α, and the tumor suppressor protein, neurofibromin (21, 22). Homozygous HAF loss is embryonic lethal in mice, suggesting a non-redundant role for HAF during development (23). Male C57Bl/6 mice heterozygous for HAF (*SART1*^*+/-*^) presented with steatosis at age 6 months that progressed to MASH and HCC without any further dietary or other intervention (23). At age >18 months, >90% of male *SART1*^+/−^ mice presented with histologically confirmed MASH-HCC (23). HAF haploinsufficiency was associated with marked upregulation of HIF-1α and increased secretion of the HIF-1α regulated chemokine, RANTES, by Kupffer cells isolated from pre-neoplastic *SART1*^+/−^ livers, implicating Kupffer cell-derived RANTES with HCC progression in the *SART1*^+/−^ mice (23). However, no HIF-1α upregulation was observed in pre-neoplastic hepatocytes from *SART1*^+/−^ mice and the cell type driving hepatocarcinogenesis associated with HAF haploinsufficiency is unclear (23).

In this study, we investigate the impact of HAF insufficiency in hepatocytes and macrophages to identify their involvement in hepatocarcinogenesis associated with HAF loss. We show that HAF depletion in hepatocytes is sufficient for MASH-HCC initiation and progression. Significantly, we reveal HAF as an essential mediator of NF-κB activity through the transcriptional regulation of TRADD and RIPK1, which has important implications in the pathogenesis of MASH and MASH-HCC.

## Materials and Methods

### Mice, cell lines and reagents

C57BL6/J mice were purchased from Jackson labs (JAX #000664) for the generation of SART1-floxed mice (see method below). The resulting mice were crossed with Alb-Cre (JAX #018961), LysM-Cre (Jax #004781) or Alf-p-Cre (generous gift from Tom Luedde at University Hospital Dusseldorf). THLE3, HepG2 and SNU475 cells were purchased from ATCC (Manassas, VA) whereas HUH7 were purchased from JCRB Cell Bank (NIBIO,Osaka, Japan). The NF-kB Luciferase Reporter HepG2 stable cell line was purchased from Signosis (Santa Clara, CA). All HepG2 and HUH7 cells were grown in Dulbecco’s modified Eagle’s medium (DMEM, ThermoFisher Scientific), while SNU475 cells were grown in RPMI-1640 (ATCC). Both types of media were supplemented with 10% FBS (Gibco) and maintained in a humidified atmosphere at 37°C in 5% CO_2_. Cell lines were authenticated by STR testing upon receipt and were routinely tested and verified as mycoplasma negative using the MycoAlert kit (Lonza, Cambridge, MA). Human recombinant TNF was purchased from BioLegend Inc (San Diego, CA), whereas MG132 and cycloheximide were purchased from Sigma-Aldrich (St Louis, MO).

### Generation of SART1 floxed mice using EASI-Crispr

Mouse embryos were injected with two preassembled guide RNA and Cas9 endonuclease ribonucleoprotein (RNP) complexes to create double stranded DNA breaks around exons 4 and 5. A long single-stranded DNA (lssDNA) donor engineered to contain a floxed version of exons 4 and 5 (grey triangles, **Fig. 1A**), flanked by 100-base pair left and right homology arms (blue) was co-injected to serve as a DNA repair template. The guide RNAs were designed to direct Cas9 mediated cutting immediately adjacent to each homology arm, which facilitated insertion of the lssDNA donor via homology directed repair (HDR). This results in replacement of thewildtype exons 4 and 5 with a floxed version. Insertion of the floxed allele was identified via simple PCR and restriction fragment length polymorphism (RFLP) analyses.

**Figure 1.**
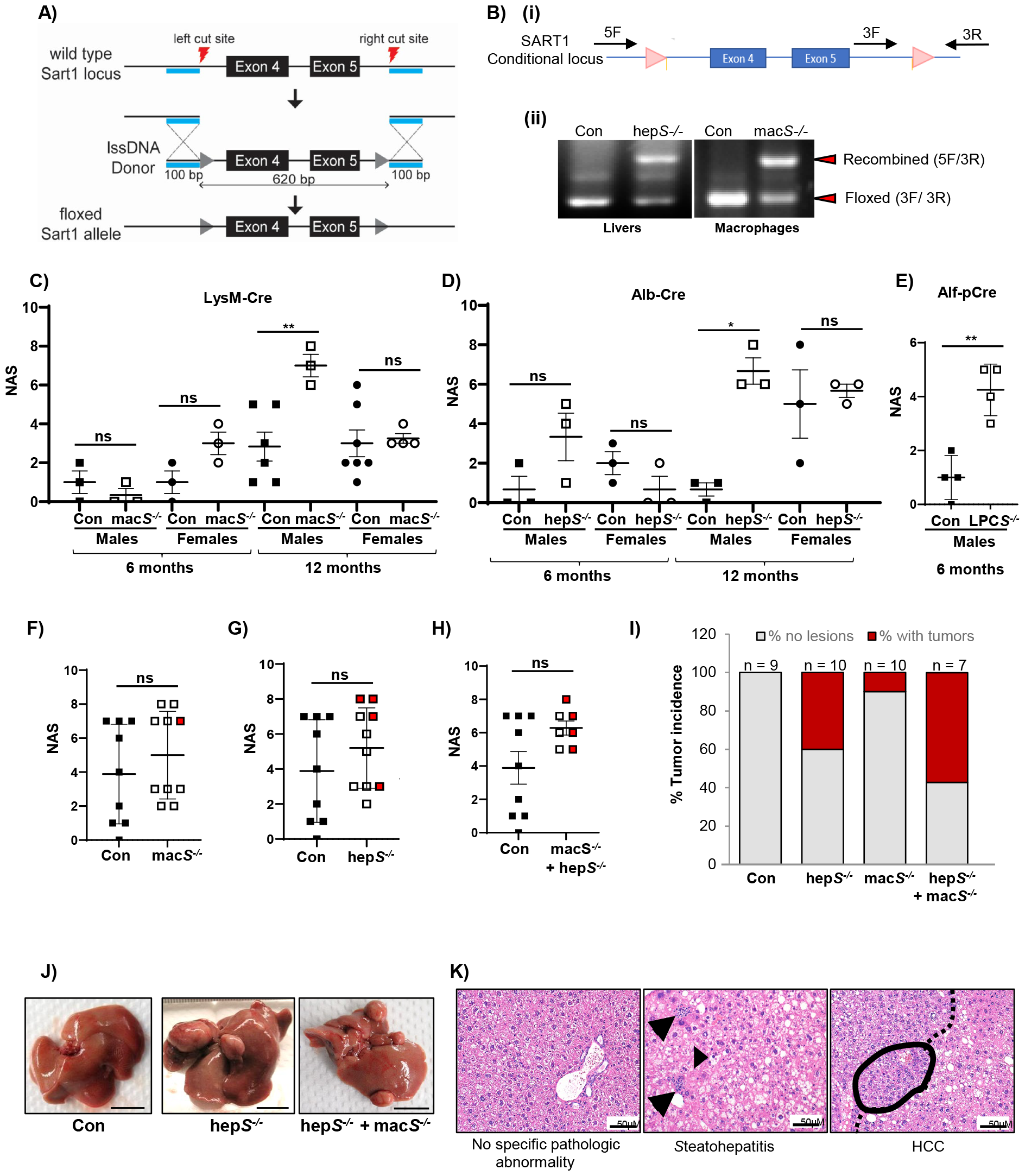
Conditional knockout of HAF in hepatocytes promotes MASH and HCC in mice. A) Easi-CRISPR method to generate floxed *SART1* allele at Exons 4 and 5. B) (i) Genotyping strategy of hepS^-/-^ and macS^-/-^ mice with primer binding sites indicated. (ii) Representative results showing floxed and recombined *SART1* alleles using isolated DNA from whole livers and isolated peritoneal macrophages from hepS^-/-^ and macS^-/-^ mice respectively. C-H) NAFLD activity score (NAS) of male and female control, hepS^-/-^, macS^-/-^ and LPC-S^+/-^ mice at age 6 and 12 months (C-E) and of male control, macS^-/-^, hepS^-/-^ or hepS^-/-^ + macS^-/-^ at 18 months (F-H) (mean ± SEM). Each shape represents NAS from an individual mouse. Red filled shapes indicate tumors histologically verified as HCC, HCA or FCA > 0.5cm. I) Quantification of tumor incidence in male control mice versus male hepS^-/-^ and macS^-/-^ mice. J) Gross appearance of livers from hepS-/- and hepS^-/-^ + macS^-/-^ mice age 18 months. Scale bar: 1 cm. K) Representative hematoxylin and eosin (H&E) stained livers from 18-month-old control or hep*S-/-* mice showing no specific pathologic abnormality (control mice) and steatohepatitis and HCC in hepS^-/-^ mice. Arrows: inflammation. Arrowhead: ballooned hepatocyte. Dotted line: LHS dysplasia. Circled: HCC. * p < 0.05; ** p < 0.01; ns: not significant

### Hepatocyte isolation

Mouse primary hepatocytes were isolated as described (24). Isolated hepatocytes were seeded at a density of 4-500,000 and 1,000,000 cells in rat tail collagen I (5 μg/cm2, Roche) pre-coated 6-well plates and T25 flasks, respectively, using DMEM low glucose containing 5% FBS and 1% of penicillin-streptomycin. After 3hrs, media was removed and hepatocytes were maintained in Williams E media supplemented with 1% glutamine and 1% penicillin-streptomycin solution. Cells were incubated at 37°C in 5% CO_2_ no more than 24h after plating.

#### DNA plasmid and siRNA transfections

For transient DNA and siRNA transfections, HepG2, HUH7, and SNU475 cells were seeded in 6-well plates, 96-well plates and chamber slides at densities of 100-150,000 cells/well, 5000 cells/well and 20,000 cells/chamber, respectively. Transfections were performed 24hrs after seeding when the cells achieved 30-40% confluency. For rescue experiments, plasmid DNA transfections were performed 24hrs after siRNA transfections (48hrs after seeding). DNA plasmids were transfected using X-tremeGENE HP (Roche Diagnostics) whereas siRNA was delivered using Lipofectamine™ RNAiMAX (ThermoFisher Scientific) according to the manufacturer’s protocols. DNA and siRNA transfected cells were incubated for 48h or 96h, respectively prior to cell lysis.

#### Immunoblotting

Cells were seeded in 6-well plates and harvested using ice-cold cell lysis buffer containing dithiothreitol and protease and phosphatase inhibitors. Protein concentration was measured using the Bradford assay. Equal amounts of protein were loaded into a Bis-Tris gel and separated by SDS-PAGE and transferred into a nitrocellulose membrane. Membranes were blocked with 5% bovine serum albumin in TBS-T, and immunoblotted. Primary rabbit polyclonal antibodies for HAF was as described (21), whereas antibodies against TRADD, RIP1, NEMO, pp65, p65, p50, IkBa, caspase-3 and -8, pRIP1, pRIP3, RIP3 were purchased from Cell Signaling Technology (Danvers, MA), antibodies against pp50 and NEMO were from Santa Cruz Biotechnology (Dallas, TX), and Bcl-xS antibody was from ThermoFisher Scientific (Waltham, MA). Imaging of the membranes was performed using the Fluorchem M imaging system (Protein Simple, San Jose, CA), and densitometry was performed using ImageJ.

### Cell viability and apoptosis assays

Cell viability of siRNA transfected cells seeded in 96 well-plates was quantified using the Resazurin cell viability assay (R&D Systems AR002). For apoptosis assays, HepG2 and HUH7 cells were seeded in 96-well and transfected with siRNA targeting HAF. For rescue experiments, cells were additionally transfected with HAF siRNA resistant DNA plasmid. 48 h post-siRNA transfection, cells were treated with the RealTime-Glo™ Annexin V Apoptosis and Necrosis Assay (Promega, Madison WI) reagents according to the manufacturer’s instructions. Depending on the experiment, cells were also treated with 20 ng/ml of TNF, ZVAD-FMK and/or nec-1 (MedChemExpress, Monmouth NJ) at the 48hr post-siRNA transfection time point. Luminescence and fluorescence were measured for a period of 48hr after adding the assay reagents.

### High-fat diet in mice

8-10 week old C57Bl/6J male mice (JAX # 000664) were fed the obesogenic Gubra Amylin NASH (GAN) diet containing 40% Kcal% fat (mostly palm oil); 22% fructose, and 2% Cholesterol (D09100301, Research diets Inc, New Brunswick NJ) with drinking water also supplemented with 23.1 g/L of D-fructose and 18.9g/L D-glucose (Sigma-Aldrich) for 7, 18, 26 and 40 weeks. Control mice were fed a normal rodent chow (#5053, Rodent diet 20, Labdiet). At the end of the dietary intervention, mice were euthanized using isoflurane and liver samples were collected and processed for histology and protein expression analysis.

### Human tissue

Human tissue microarrays containing de-identified patient liver samples were purchased from Protein Biotechnologies (TMA1427/1428) and stained using a previously validated monoclonal HAF antibody as previously described (25). Slides were scanned using the Aperio AT2 digital scanning system and HAF nuclear staining intensity quantitation was performed using Aperio digital imaging software (Leica Biosystems, Buffalo Grove IL) and statistical analysis performed using unpaired Student’s T-tests in GraphPad Prism software 10.10.2 (GraphPad Software, San Diego CA).

All other methods are described in supplemental methods.

## Results

### Hepatocyte-specific HAF deletion promotes tumor formation in both male and female mice

To investigate the impact of conditional HAF deletion on hepatocarcinogenesis, we generated C57BL/6 mice bearing flox sites flanking exons 4 and 5 of the *SART1* gene using Easi-CRISPR (Efficient additions with ssDNA inserts-CRISPR) (**Fig. 1A**) (26). Mice homozygous for the floxed allele (*SART1*^*FL/FL*^) were crossed with mice expressing Cre recombinase under the control of the mouse albumin enhancer and promoter as well as the mouse alpha-fetoprotein enhancers (Alfp-Cre) to generate liver parenchymal cell (LPC)-specific *SART1*-knockout mice (27). No Cre-expressing *SART1*^*FL/FL*^ pups were obtained suggesting embryonic lethality reminiscent of that seen in mice with constitutive deletion of *SART1* in all body cells at E11.5 (23). However, Alfp-Cre mice heterozygous for *SART1* (Alfp-cre^+/-^, SART1^+/FL^, LPC-S^+/-^) were viable. We also crossed *SART1*^*FL/FL*^ mice with those expressing Cre recombinase driven by the mouse albumin promoter (Alb-Cre; *Alb-Cre*^*+/-*^ *SART1*^*FL/FL;*^ *hepS*^*-/-*^) or lysozyme M (Lys-M-Cre; *LysM-Cre*^*+/-*^ *SART1*^*FL/FL;*^ *macS*^*-/-*^). These mice survived to adulthood and showed > 50% recombination of *SART1* detected by PCR in the livers or isolated macrophages from these mice compared to *Cre*^*-/-*^ *SART1*^*FL/FL*^ mice (control) and significant reduction in *SART1* transcript detected by qPCR (**Fig. 1B, S1A**). H&E-stained liver sections of mice at various ages were examined by pathologist (KE) and assigned a NAFLD-activity score (NAS) based on established criteria including the prevalence of macro- and micro-vesicular and micro-vesicular steatosis, hepatocellular hypertrophy and inflammation (**Fig. 1C-E**) (28). The pathologist was blinded to the genotypes of the mice during analysis. We observed significantly higher NAS in male *mac*S^-/-^ and *hepS*^*-/-*^ mice at 1 year and in male *LPC*-S^+/-^ mice at age 6 months compared to age-matched controls (**Fig 1C-E**). This was accompanied by elevation of serum transaminase, ALT in both male and female *hepS*^*-/-*^ mice compared to controls from age 6 months indicating hepatocyte injury (**S1B**). At age 18 months, NAS were not significantly different between *mac*S^-/-^, *hep*S^-/-^ or *mac*S^-/-^ + *hep*S^-/-^ mice compared to age-matched control (**Fig. 1F-H, S1C**). However, we detected liver tumors (histologically confirmed as HCC, HCA or FCA > 0.5mm) in 42% (5 of 12) and 25% (3 of 12) of 18-month-old male and female *hep*S^*-/-*^ mice respectively and with histologically confirmed HCC in 33% (4 of 12) and 17% (2 of 12) of male and female *hep*S^*-/-*^ mice respectively (**Fig. 1G, S1D**). By contrast, *SART1* deletion in macrophages resulted in liver tumors in only 1 of 10 male mice at age 18 months, which was histologically confirmed as HCA (**Fig. 1F**) and not in female mice (not shown). Intriguingly, the combination of *SART1* deficiency in both macrophages and hepatocytes resulted in potentially additive effects with 57% tumor incidence (4 of 7; **Fig. 1H-I**). Representative images of gross liver tumors and liver histology are shown in **Fig 1J-K and S1E-F**. Thus, we show that hepatocyte-specific loss of HAF induces tumors in both male and female mice, whereas macrophage-specific loss only induces tumors at low penetrance in male mice. The combination of both hepatocyte- and macrophage-specific loss of HAF contributes an additive effect to tumor incidence.

### Hepatocyte-specific HAF deletion is associated with lipid dysregulation

To characterize the metabolic changes associated with progression to HCC, we performed lipidomics profiling of livers from male control and hepS^-/-^ mice at ages 6, 12 and 18 months (5 mice per group) as well as liver tumors from three male 18-month-old hepS^-/-^ mice. The lipid species identified from the livers or liver tumors from hepS^-/-^ mice were compared to their appropriate age-matched control mice. Using the LipidR software tool to implement lipid set enrichment analysis, we found significant enrichment of triglycerides (TG) in hepS^-/-^ mice at 6 and 12 months of age (**Fig. 2A-B**), consistent with the increased lipid accumulation and NAS observed in the livers of these mice (**Fig. 1D**) (29). At age 18 months, the tumor bearing livers of the hepS^-/-^ mice showed profound lipid dysregulation with significant changes in multiple lipid classes (**Fig. 2C**). In this regard, the acylcarnitines (ACars), particularly long chain ACars (> 18 carbon atoms) and TG were the most highly upregulated detected lipid classes within the hepS^-/-^ tumors compared to livers from age matched wild-type littermates (**Fig. 2C-E**), with the most profound upregulation observed in ACar C20:1, which was also significantly upregulated in livers of 6-month-old HepS-/- mice compared to age matched wild-type littermates (**S2A**). This suggests that the accumulation of TG and ACars are early events in the metabolic dysfunction associated with HAF loss. Since long chain fatty acids must be esterified to ACars to enter the mitochondria for fatty acid oxidation (FAO), the accumulation of long chain ACars is consistent with the defective FAO that we previously reported in the *SART1*^*+/-*^ mice (23, 30) and may also indicate mitochondrial dysfunction and/or a metabolic shift to lipid oxidation (30). In addition to ACar upregulation, we observed increases in C16:0 and C24:1 ceramides (Cers) in the hepS-/- tumors compared to normal liver from non-tumor bearing hepS-/- mice and wild-type control mice (**S2B**). In mice, upregulation of C16:0 Cer in the liver is associated with altered glucose and lipid metabolism, increased oxidative stress, enhanced susceptibility to apoptosis and HCC (31-36). In this regard, both <u>ACars</u> and Cers can cause increased cell death and/or reactive oxygen species (ROS) that drive inflammation, in part through activation of the NF-κB pathway, thus driving the lipotoxicity associated with MASH (37-40).

**Figure 2.**
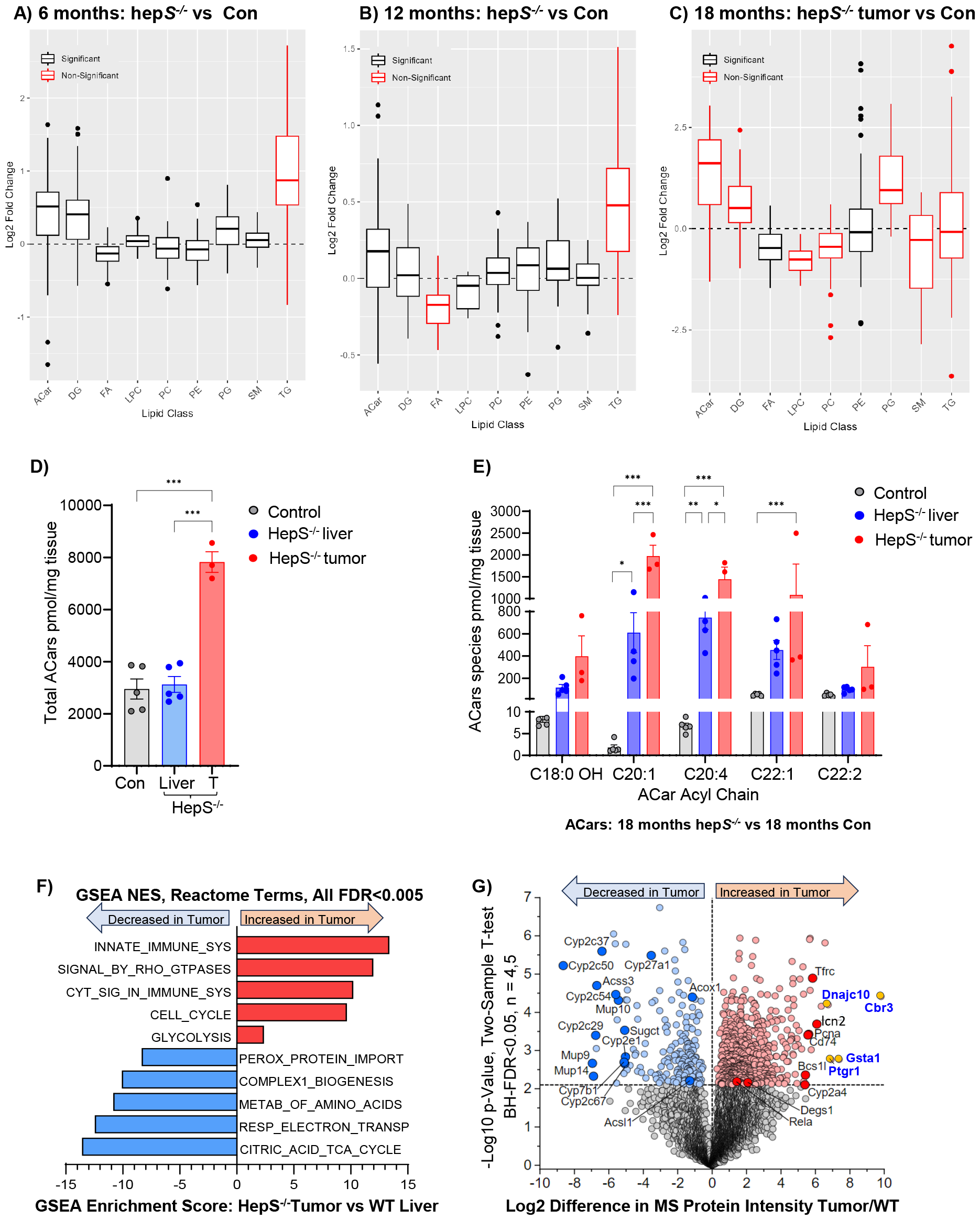
Inflammatory and metabolic dysfunction are associated with progression to HCC. A-C) Box plots showing changes in indicated lipid species in livers of male *hepS*^*-/-*^ vs control mice at ages 6, 12 and 18 months (5 mice/group) or liver tumors of 18-month-old male *hepS*^*-/-*^ mice (3 mice). D-E) Levels of total acylcarnitines (ACars, D), and most abundant ACars species, (E), in non-tumor bearing livers (L) or liver tumors (T) from 18-month old male control (L only) or hepS^-/-^ (L and T) mice. F) Normalized enrichment scores (NES) obtained through GSEA of differentially expressed proteins from liver tumors of male 18-month-old hepS-/- mice (4 mice) compared to age matched control mice (5 mice). G) Volcano plot showing significantly upregulated and downregulated proteins in hepS-/- tumors compared to age-matched control littermates (two sample t-Test with Benjamini-Hochberg (BH)-correction, FDR = 0.05, n = 4,5). Yellow circles: proteins involved in antioxidant response. Red circles: Proteins involved in proliferation/inflammation. Blue circles: Downregulated metabolic proteins.

To investigate the pathway perturbations associated with liver tumor development in the HepS^-/-^ mice, we applied mass spectrometry (MS)-based global proteomics with label-free quantification to compare the proteome of tumors from male 18-month-old *hepS*^*-/-*^ mice (n=4) to that of age-matched control littermates (n=5) (41). This analysis quantified the expression of 3883 proteins. Differential expression analysis between hepS^-/-^ and control livers followed by Gene Set Enrichment Analysis (GSEA) (42, 43) with Reactome pathway terms (44) revealed that the top upregulated pathways in the *hepS*^*-/-*^ tumors included those involved in inflammatory activation (Innate immune system/cytokine signaling), proliferation, migration, invasion, metastasis (activation of Rho GTPases, cell cycle) and a shift towards glycolysis (**Fig. 2F**), all of which are well-validated characteristics of HCC ^66^. By contrast, the top downregulated pathways in the *hepS*^*-/-*^ tumors were remarkably enriched for those involved in lipid metabolism and FAO (**Fig. 2F**). Analyzing differentially expressed proteins in our global proteomics data revealed significant upregulation of key proteins linked to the antioxidant response (Cbr3, Dnajc10, Gsta1, Ptgr1), proliferation (Pcna, Lcn2, Tfrc) inflammation (RelA, Cd74) and sphingolipid metabolism (Degs1) in the *hepS*^*-/-*^ tumors (**Fig. 2G**). Intriguingly, the majority of upregulated proteins linked to the antioxidant response are regulated by the transcription factor, Nrf2, which was not detected in this screen, but which is itself regulated by NF-κB and cooperates with the latter to protect hepatocytes from oxidative stress (45-49). Indeed, RelA (p65), the key transcriptional subunit of the NF-κB heterodimer was significantly upregulated in the *hepS*^*-/-*^ tumors (**Fig. 2G**), implicating NF-κB activation as a central mediator of the proteome changes associated with the hepS^-/-^ tumors. The top downregulated proteins in the *hepS*^*-/-*^ tumors were primarily the cytochrome P450 proteins, which have diverse roles in liver metabolism (**Fig. 2G**). These proteins, together with Acsl1, a protein which catalyzes one of the first steps of FAO, were also significantly downregulated at both the transcript and protein levels in the *SART1*^*+/-*^ tumors, consistent with the decreased FAO that we have previously reported (23). We also observed significant downregulation of various isoforms of major urinary proteins (MUPs) in the *hepS*^*-/-*^ tumors (**Fig. 2G**), whose deficiency has been associated with obesity and type 2 diabetes in mice (50). Taken together, the data suggest that *hepS*^*-/-*^ tumors express protein expression changes consistent with increased inflammation/proliferation/resistance to oxidative stress and decreased FAO, the former driven in part by NF-κB activation, whereas the latter driven by deficiencies in Cyp450 expression.

### Hepatocyte-specific HAF deletion is associated with downregulation of the NF-κB pathway

To investigate the potential association of HAF with the NF-κB pathway, we investigated the phosphorylation of the transcriptional subunits of NF-κB: p65 (RelA) and p50. Phosphorylation of p65 at S536 (P-p65) promotes p65 nuclear translocation and transactivation, whereas phosphorylation of p50 at S337 (P-p50) is required for its DNA binding activity (51-53). We observed that HAF depletion in both mouse models was associated with significant and profound reduction in p50 and p65 phosphorylation (**Fig. 3A-C, S3A**). To investigate the effect of HAF on the NF-κB pathway independent of any potential liver pathology in the conditional knockout mice, we performed HAF siRNA knockdown in a panel of liver cancer cell lines. Indeed, we show that HAF knockdown using two independent siRNAs resulted in a marked reduction in levels of P-p65 and P-p50 as well as in the upstream components of the NF-κB pathway, TRADD, RIPK1 and NEMO in both HEPG2 and SNU475 cells (**Fig. 3D, S3B**,**D**). We also confirmed reduction in total and nuclear P-p65 through immunocytochemistry of SNU475 cells (**S3C**) and show that the reduction in P-p65 and TRADD occurred as soon as 48 hours after HAF siRNA transfection (**S3D**). Furthermore, HAF siRNA profoundly attenuated the TNF-induced increase in P-p65/P-p50 and in the luciferase activity of cells stably expressing the NF-κB response element (RE) fused to a luciferase reporter (**Fig. 3D-E**). Conversely, HAF overexpression further enhanced TNF-induced levels of P-p65/P-p50 and of NF-κB RE reporter activity (**Fig. 3F-G**). We also observed more subtle increases in TNF-induced levels of TRADD, RIPK1 and NEMO in HAF-overexpressing cells (**Fig. 3F**). Taken together, our data suggests that HAF is required for the phosphorylation of p65 and p50, and is both necessary and sufficient for TNF-induced NF-κB transcriptional activity.

**Figure 3.**
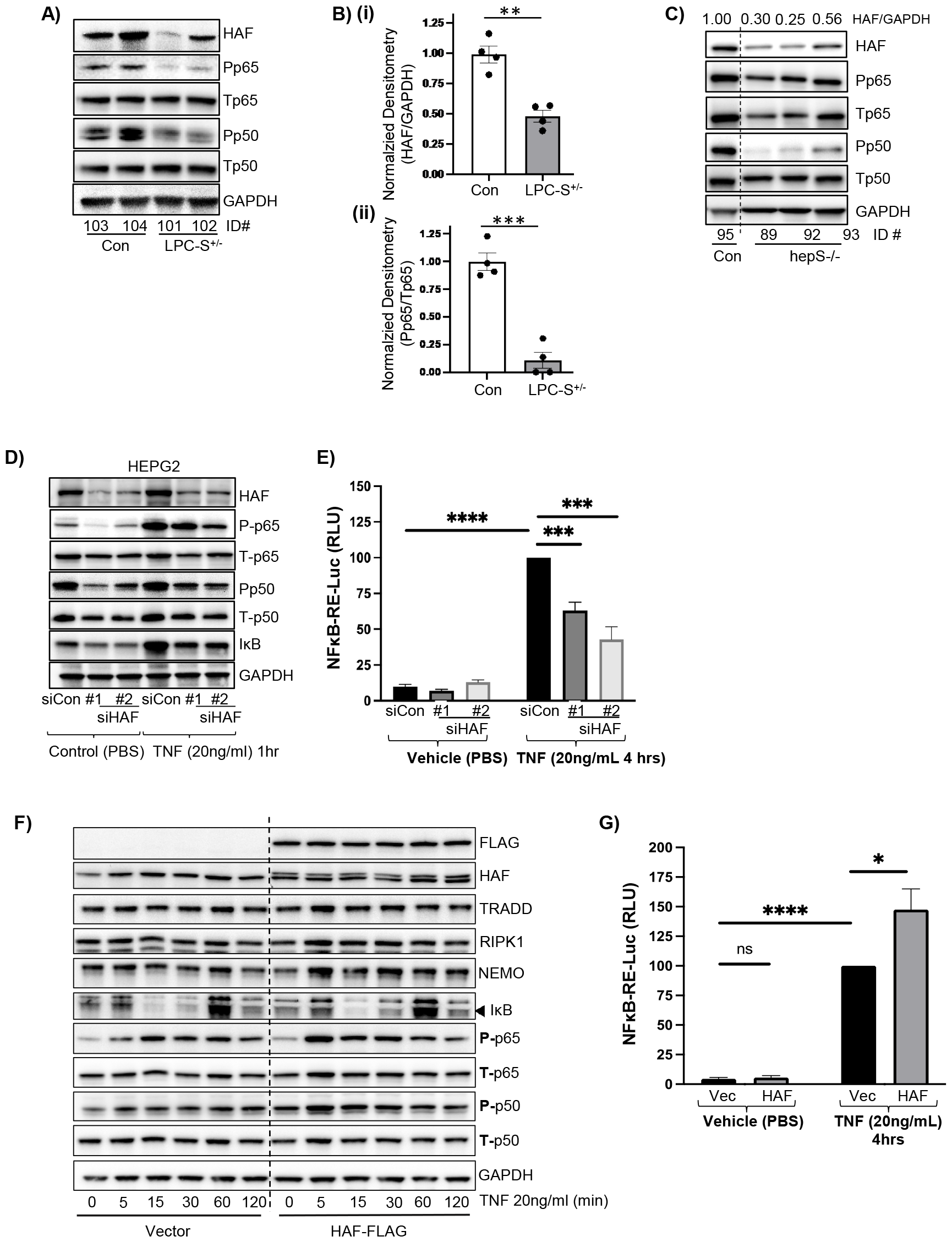
HAF depletion *in vivo* and *in vitro* is associated with decreased NF-κB activity. A-B) Effect of HAF depletion on phosphorylation of NF-κB p65 and NF-κB p50 via western blotting in whole liver lysates from 6-month-old LPC-S^+/-^ mice (A), with densitometric quantitation of HAF/GAPDH and P-p65/total p65 in B (i) and (ii) respectively; and in isolated hepatocytes from 3-month-old male hepS^-/-^ mice (C), with quantitation shown above plots. Each lane or data point on plots represents data from an individual mouse. Data are representative of at least 2 independent experiments. D-E) Effect of transient transfection with two independent HAF siRNAs versus non-targeting control siRNA on (D), protein levels of P-p65/T-p65 and P-p50/T-p50, or (E) NF-κB response element luciferase (NF-κB-RE Luc) reporter activity in HepG2 cells (representative data from 3 independent experiments). F) Western blot showing effect of HAF overexpression on indicated NF-kB pathway components after treatment with indicated durations of 20ng/ml TNF in HepG2 cells G) Effect of HAF overexpression on NF-κB-RE Luc in the absence of presence of TNF. Data are representative of three independent experiments and are shown as mean ± SEM. * p < 0.05; ** p < 0.01; *** p < 0.001; **** p < 0.0001 ns: not significant

### HAF promotes transcriptional activation of TRADD and RIPK1

We next sought to elucidate the mechanism by which HAF regulates NF-kB activation. As previously observed, HAF siRNA markedly depleted many components of the canonical NF-kB signaling cascade including TRADD, RIPK1, TAK1 and NEMO as well as P-p50 and P-p65 (**Fig. 4A**). These proteins were similarly reduced in isolated hepatocytes from LPC-S^+/-^ mice (**S4A**). Subsequent studies to identify potential mechanism(s) of HAF-mediated NF-κB regulation focused on these 4 proteins since they function upstream of p50 and p65. We found that treatment with the proteasome inhibitor, MG132, while attenuating the siHAF-induced decrease in NEMO, did not ameliorate decreases in TRADD or RIPK1 suggesting that HAF knockdown may promote the degradation of NEMO, but not of TRADD and RIPK1 (**Fig. 4B**). RIPK1 has also been reported to be a substrate of lysosomal degradation (54). However, co-treatment of HAF siRNA-transfected cells with the lysosomal inhibitor, chloroquine, only partially attenuated the siHAF-induced decrease in RIPK1 but did not affect the siHAF-induced decrease in TRADD, TAK1 or NEMO (**S4C**). Additionally, although co-treatment of HAF siRNA-transfected cells with cycloheximide (inhibitor of mRNA translation) resulted in a trend of decreased stability of these proteins (particularly TAK1 and NEMO), these changes were not significant suggesting that decreased protein stability is not the primary mechanism driving their decrease in the context of HAF siRNA (**Fig. 4C, S4B**). Consequently, we next investigated the impact of HAF on the transcription of *TRADD, RIPK1, TAK1* and *NEMO*. Intriguingly, qRT-PCR analysis demonstrated that *RIPK1* and *TRADD* mRNA levels were significantly reduced with HAF siRNA, whereas no significant changes were observed in the transcript levels of *TAK1* and *NEMO* (**Fig. 4D, S3D**). The transcriptional decreases in *TRADD* and *RIPK1* were observed across multiple exon boundaries, including previously reported alternative exons, suggesting that HAF knockdown decreases overall transcription of TRADD and RIPK1 mRNA and does not promote their alternative splicing (**Fig. S4F-G**) (55). Conversely, HAF overexpression in HEPG2 cells significantly induced RIPK1 and TRADD transcription suggesting HAF regulates the transcription of *TRADD* and *RIPK1* (**Fig 4E**).

**Figure 4.**
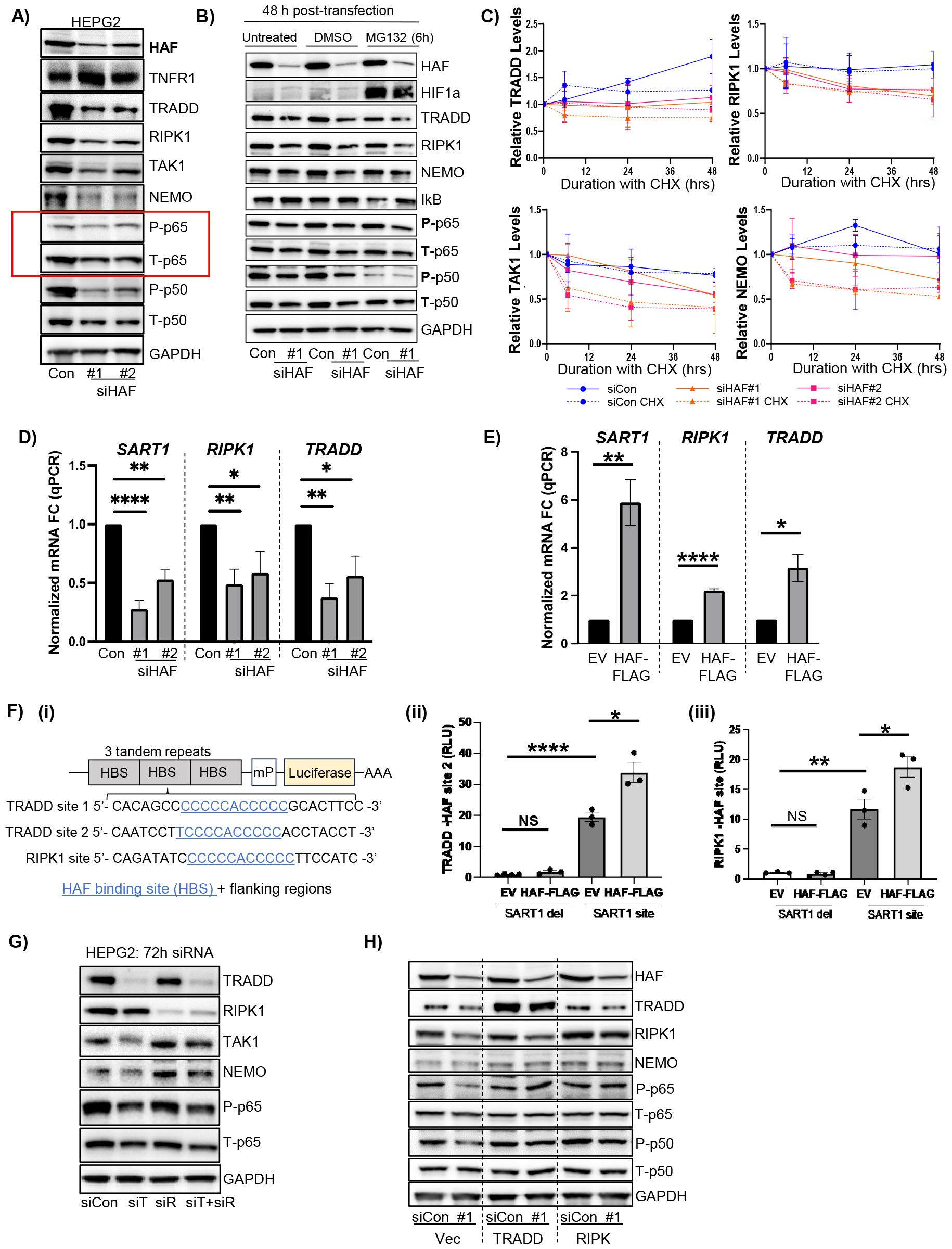
HAF regulates transcription of *TRADD* and *RIPK1*. A) Western blot showing effect of HAF siRNA on major components of the canonical NF-kB pathway in HepG2 cells. B) Impact of 20μM MG132 (6hrs) treatment on major NF-kB pathway components impacted by HAF siRNA. C) Quantitation of relative levels of TRADD, RIPK1, TAK1 and NEMO protein normalized to GAPDH after treatment with 25 ug/ml of CHX 24hrs post HAF siRNA transfection in HepG2 cells, and lysed at 30, 48 or 72hrs based on average values from western blots shown in S4B. Data are the mean of 3 independent experiments ± SEM. D) Quantitative RT-PCR analysis of *SART1, RIPK1* and *TRADD* mRNA levels after transfection with HAF siRNA (D), or stable HAF overexpression (E), in HepG2 cells. Data is representative of 3 independent experiments. F) (i) Construction of luciferase reporters containing 3 tandem repeats of the HAF binding site (HBS) and flanking regions (SART1 site) or those bearing only the flanking regions without the HBS (SART1del). (ii-iii) Relative luciferase activity of TRADD site 2 (ii) and RIPK1 site (iii) in control and HAF overexpressing ACHN cells. Data are representative of 3 independent experiments (mean± SEM). G) Western blot showing the impact of TRADD and RIPK1 siRNA on components of the NF-kB pathway in HepG2 cells. H) Impact of overexpression of TRADD or RIPK1 on P-p65/T-p65 and P-p50/T-p50 in HEPG2 cells transfected with HAF siRNA.

HAF was initially identified as a DNA-binding protein that regulates the hypoxia-transcription of the *EPO* gene (19). We previously confirmed the binding of HAF to its consensus DNA binding site (CCCCRRCCCC) within regulatory regions of *VEGFA* and *OCT-3/4* to contribute to their hypoxia-induced transcription (20). To determine whether HAF also regulates transcription of *TRADD* and *RIPK1*, we searched the regulatory regions of *TRADD* and *RIPK1* for potential HAF binding sites (HBS). We identified two potential binding sites flanking the *TRADD* transcriptional start site (TSS) based on TRADD-201 transcript ENST00000345057: Site 1 was located 6604 bases upstream of the TSS, whereas Site 2 was located 1319 bases downstream of the TSS. Similarly, we identified a single HBS within the reverse motif 800 base pairs upstream of the TSS of RIPK1 (based on ENST00000259808) (**Fig. 4Fi**). To investigate the functionality of these putative HBS’s, we generated 3 tandem repeats of the HBS and flanking regions for each putative HBS (SART1 site) as well as mutant versions that contained the flanking regions but excluded the HBS (SART1 del) in tandem with a luciferase reporter construct driven by a minimal promoter (**Fig. 4Fi**). The presence of an HBS on the TRADD site 1 construct did not promote an increase in luciferase activity (**S4E**) suggesting that this HBS is not transcriptionally active. However, the presence of the HBS from TRADD site 2 and the RIPK1 site significantly increased luciferase reporter activity suggesting that these HBS’s convey transcription activation (**Fig. 4Fii-iii**). Furthermore, transfection of these reporters into cells overexpressing HAF resulted in significant upregulation of luciferase activity compared to empty vector expressing cells, suggesting that these HBS’s are active and convey HAF-driven transcriptional activity (**Fig. 4Fii-iii**). By contrast, no effect of HAF overexpression was observed in the reporter constructs that lacked the HBS. To determine whether the reduction in *TRADD* and *RIPK1* were the primary drivers of reduced P-p65 and P-p50 associated with HAF loss, we investigated whether siRNA knockdown of TRADD and RIPK1 could recapitulate the effects of HAF loss on components of the NF-κB pathway. Indeed, we found that knockdown of TRADD (but not RIPK1) reduced levels of TAK1, NEMO and P-p65 in HEPG2 cells (**Fig. 4G**). Furthermore, overexpression of either TRADD or RIPK1 was sufficient to rescue the decrease in P-p65 and P-p50 associated with HAF siRNA (**Fig. 4H**). Thus, we show that HAF is necessary and sufficient for the transcription of *RIPK1* and *TRADD* through the presence of HBS’s in the regulatory regions of these genes. Loss of HAF results in decreased transcription of *RIPK1* and *TRADD* resulting in decreased phosphorylation of p65 and p50.

### HAF loss promotes spontaneous apoptosis

It has been shown that ablation of NF-κB pathway components (p65, NEMO, TAK1) leads to liver damage and HCC in mice due to increased cell death (56-60). To investigate the impact of HAF on cell death, we examined the impact of HAF downregulation on cell viability. We found that HAF knockdown using two independent siRNAs significantly reduced the viability of HEPG2 and PLC/PRF5 HCC cells, as well as THLE3 immortalized human hepatocytes (**Fig. 5A, S5A, D**). Moreover, HAF knockdown also resulted in reduced RIPK3 phosphorylation and spontaneous cleavage of caspase 8 and 3 that was clearly apparent at 72hrs post siRNA transfection (**Fig. 5B, S5C**). HAF siRNA also resulted in a profound change in the splicing of the anti-apoptotic protein, Bcl-X_L_ to favor its pro-apoptotic form - Bcl-X_s_ (**Fig. 5C, S5E**) which may reflect HAF’s previously reported role in the regulation of mRNA splicing (61, 62). Conversely, overexpression of HAF attenuated the TNF-induced decrease in Bcl-X_L_/Bcl-X_s_ ratio (**5D**). Furthermore, assessment of apoptosis in real time showed an increase in Annexin V positivity in cells transfected with HAF siRNA (**Fig. 5E, S5B**,**F**), confirming apoptosis. Since the increase in Annexin V positivity, while suggesting death via apoptosis, does not necessarily preclude death via necroptosis, we treated HAF siRNA-treated cells with the pan-caspase inhibitor Z-VAD-FMK, which blocks apoptosis, or the RIPK1 inhibitor, Necrostatin-1, which blocks necroptosis. Treatment with ZVAD-FMK but not necrostatin-1 protected against cell death (**Fig. 5E**) suggesting that cell death associated with HAF siRNA was primarily due to increased apoptosis. Additionally, treatment with ZVAD-FMK profoundly decreased levels of cleaved caspase 3 in cells transfected with HAF siRNA (**Fig. 5F**), confirming apoptosis as the major mechanism of cell death induced by HAF siRNA. We noted that ZVAD-FMK treatment was associated with increased levels of NEMO in cells transfected with siHAF suggesting that the siHAF-induced decrease in NEMO may be due to increased apoptosis associated with siHAF and may thus be an indirect effect of HAF knockdown (**Fig. 5F**). TNF stimulation of cells potentiated the cell death response associated with HAF siRNA (**S5D, F**) suggesting that HAF protects against TNF-induced apooptosis. To confirm that the increase in spontaneous apoptosis was indeed mediated by HAF deficiency, we co-transfected cells with HAF siRNA and a HAF construct resistant to siRNA knockdown, FLAG-HAF(r). Overexpression of FLAG-HAF(r) markedly decreased the cleavage of caspase 8 and 3, and significantly decreased annexin V staining associated with siHAF, confirming that HAF loss indeed promotes spontaneous apoptosis (**Fig. 5G-H)**. Consistent with our *in vitro* data, immunohistochemical analysis of liver sections from 6-month-old hepS^-/-^ and LPC-S^+/-^ mice revealed significant increases in cleaved caspase 3 (**Fig. 5I-J**), and in significant decreases in the ratio of Bcl-X_L_ to Bcl-X_S_ (**S5G-H**) suggesting that HAF deficiency results in significantly increased apoptosis *in vivo* that may contribute to the development of MASH and MASH-HCC in these mice. The increase in spontaneous apoptosis was confirmed by increased Annexin V staining in primary hepatocytes isolated from 5-month-old male LPC-S^+/-^ mice (**S5I**). Thus, collectively, our *in vitro* and *in vivo* data suggest that HAF downregulation promotes spontaneous apoptosis and potentiates apoptosis induced by TNF.

**Figure 5.**
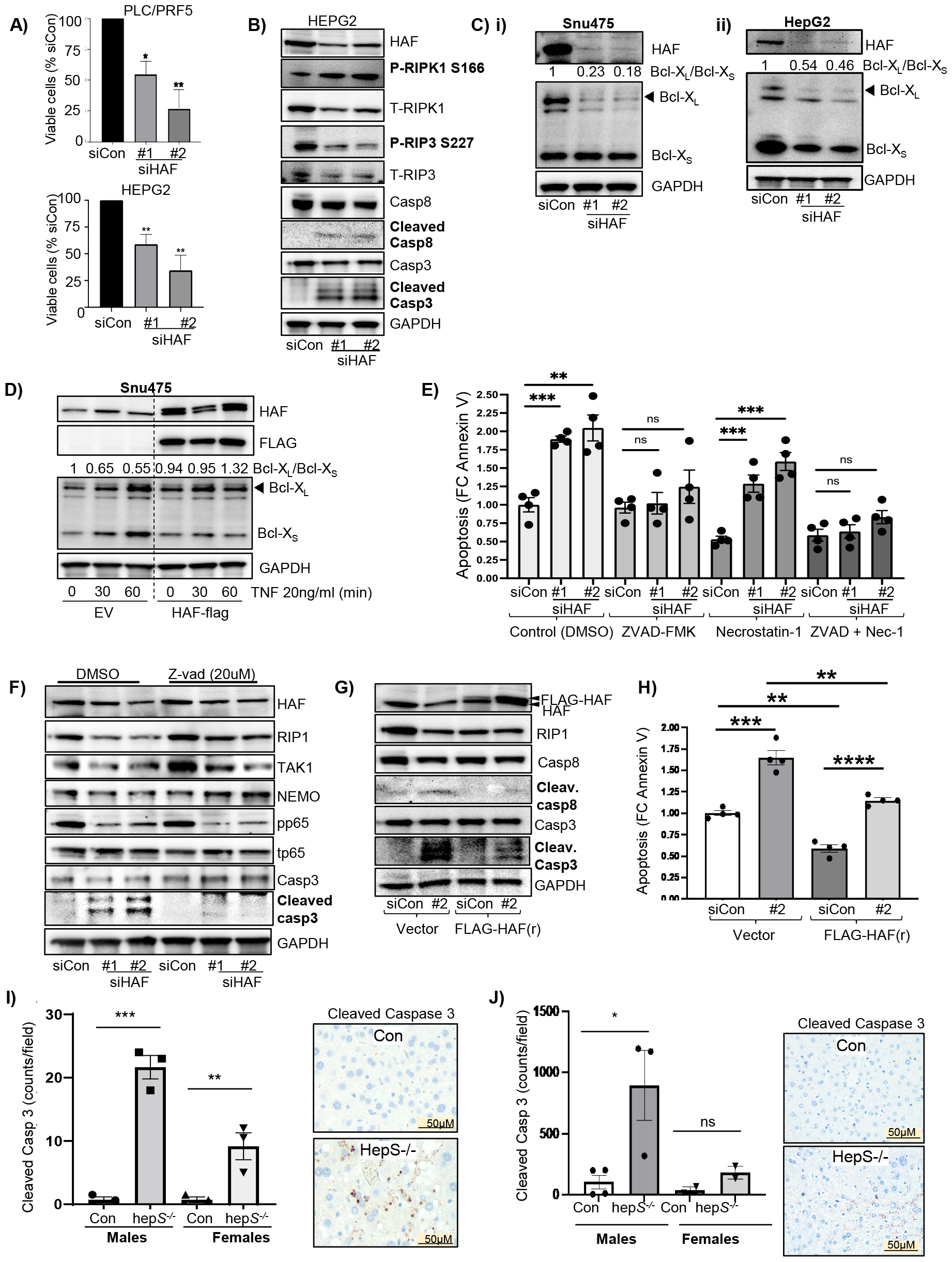
HAF depletion triggers spontaneous apoptotic cell death. A) Impact of HAF siRNA on cell viability (resazurin assay) in indicated cell lines. B) Western blot showing upregulation on cell death markers (B) and on splicing of Bcl-X_L_ and Bcl-X_s_ (C) in indicated cell lines transfected with HAF siRNA. D) Effect of HAF overexpression on splicing of Bcl-X_L_/Bcl-X_s_ after treatment with TNF. E) Effect of HAF siRNA ± the pan-caspase inhibitor ZVAD-FMK and/or necroptosis/RIPK1 inhibitor, necrostatin-1 on annexin V staining in HepG2 cells. Inhibitors were added 48 hrs after siRNA transfection. F-G) Western blots showing effects of (F) HAF siRNA ± ZVAD-FMK for 72hrs, or (G) overexpression of a HAF siRNA-resistant construct (FLAG-HAF-r) in the context of HAF siRNA in HEPG2 cells. H) Impact of FLAG-HAF-(r) in the context of HAF siRNA on annexin V staining in HEPG2 cells. I-J) Quantitation of immunohistochemical staining for cleaved caspase 3 in livers of 6-month-old male and female hepS^-/-^ (I) and LPC-S^+/-^ mice (J).

#### Exposure to a HFD suppresses HAF, RIPK1 and TRADD

Our data thus far suggest an essential role for HAF in the activation of the NF-κB pathway, mediated primarily through HAF’s role in regulating transcription of *TRADD* and *RIPK1*. To investigate potential physiological regulators of HAF, we investigated the impact of hypoxia, or TNF, conditions prevalent during hepatic steatosis and MASH, on HAF levels. Exposure of immortalized hepatocytes (THLE3 cells) or HCC cells to 4hrs hypoxia profoundly decreased HAF levels (**Fig. 6A**), which is in agreement with our previous studies showing that HAF is modulated by hypoxia intensities and duration (63). Exposure of primary hepatocytes isolated from control or LPC-S^+/-^ mice to hypoxia also resulted in substantial decreases in HAF, which was associated with striking increases in HIF-1α (**Fig. 6B**), consistent with HAFs previously reported role as an E3 ligase for HIF-1α (21). Conversely, treatment of primary hepatocytes with TNF resulted in significant induction of HAF (**Fig. 6C**), which is consistent with our proposed role for HAF as a critical component of the NF-κB response to TNF. To determine the involvement of HAF in progression to HCC, we examined the impact of long-term feeding of the Gubra-Amylin NASH (GAN) high-fat diet (HFD) on HAF levels. Livers of male C57BL/6 mice fed the GAN HFD diet for 38 weeks show histological hallmarks of human MASH including steatosis, inflammation, hepatocyte ballooning degeneration and fibrosis (64). Feeding of the HFD diet for 7 weeks or more resulted in hepatic lipid accumulation and elevated NAS (as expected) (**Fig. 6D**). Strikingly, livers of mice fed the HFD diet for 7-26 weeks showed profound suppression of HAF, TRADD and P-p65 compared to mice fed a chow diet, which were then strongly induced after 40 weeks on HFD diet (**Fig. 6E**). By contrast, no induction of HAF was seen in livers of mice fed a chow diet for 48-72 weeks compared to 24 weeks suggesting that the HAF increase seen with mice fed the HFD diet for 40 weeks was not due to simple aging of the mice (**S6**). Significantly, we observed a similar pattern of HAF expression in human livers using tissue microarrays whereby HAF was significantly decreased in steatotic human livers compared to normal human livers but then progressively increased from simple steatosis to cirrhosis through HCC, where HAF levels were significantly higher compared to normal liver (**Fig. 6F**). Taken together, the data suggest that HAF is suppressed during simple steatosis (possibly due to hypoxia and other cellular mediators), which exacerbates hepatocyte apoptosis during cellular stress that promotes progression to HCC suggesting a tumor suppressor role. Paradoxically, HAF is strongly induced during steatohepatitis/MASH/cirrhosis, in part due to elevated TNF signaling, which is concomitant with activation of the NF-κB pathway.

**Figure 6.**
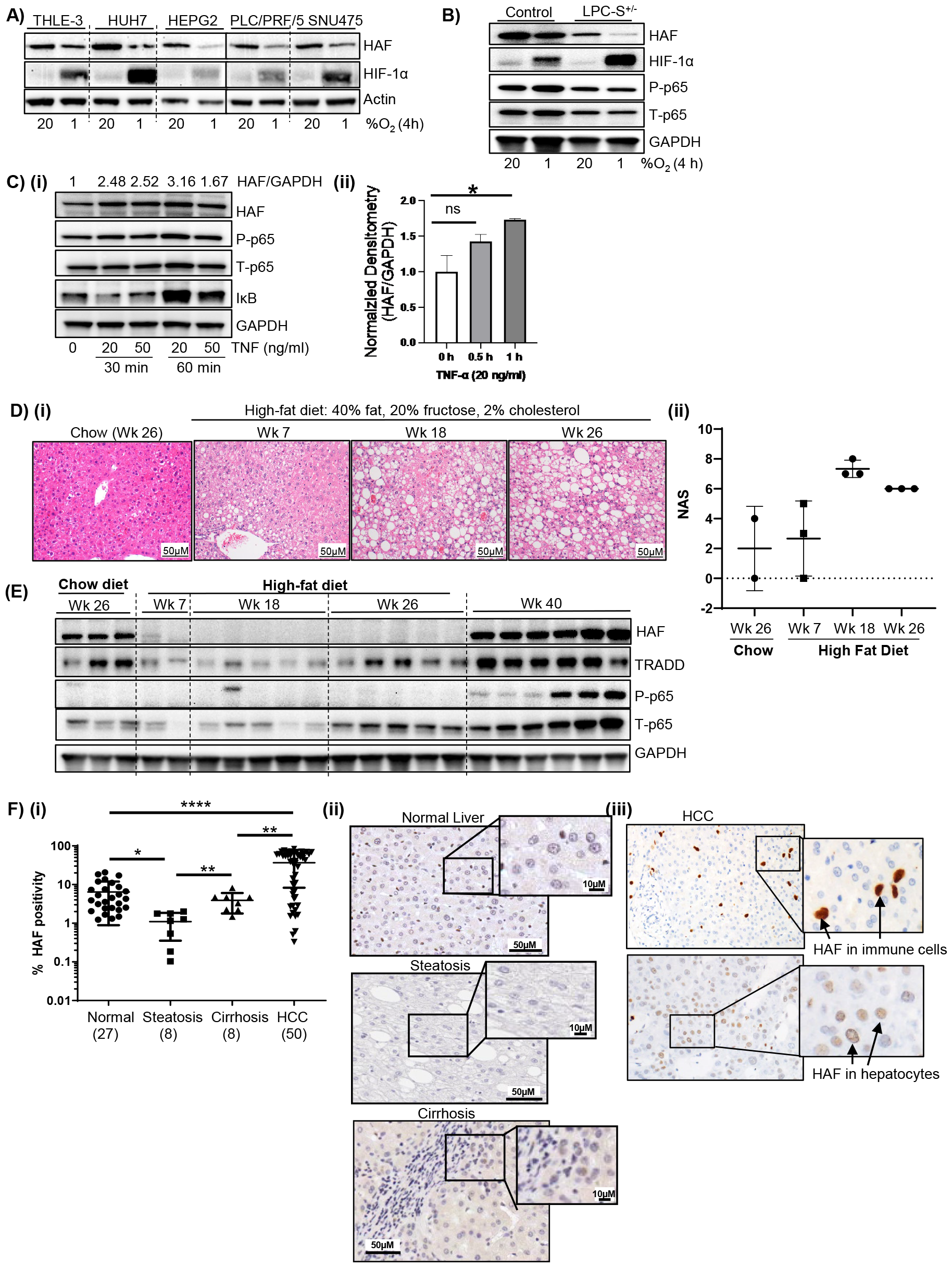
HAF is decreased during progression to MASH and HCC in mice and humans. A-B) Western blots showing the impact of hypoxia (1% O_2_, 4 hrs) on indicated cell lines (A), and primary hepatocytes isolated from 3-month-old control and LPC-S+/- mice (B). C) Western blots showing effect of TNF on HAF and NF-κB pathway components (i) with quantitation in (ii). D-E) H&E (D(i)), NAS (D(ii)) and western blots (E), from livers of C57BL/6 mice fed a high fat diet and sugar water for 7, 18 or 26 weeks compared to mice fed with a chow diet for 26 weeks. F) (i) Percentage of HAF positivity in human liver sections using tissue microarrays quantitated using Aperio digital quantitation. Representative images of HAF staining in normal and various non-HCC pathology (ii) and HCC (iii) are depicted.

## Discussion

In this study, we reveal a novel and essential role for HAF in the canonical NF-κB pathway through the transcriptional regulation of TRADD and RIPK1. Although our previous studies using global HAF/SART1 haploinsufficiency in mice implicated Kuffer-cell associated RANTES as a key mediator of HCC progression associated with HAF insufficiency (23), here we show that HAF loss in hepatocytes (but not in macrophages) is sufficient to induce HCC in mice (**Fig. 1I**). Furthermore, we show that HAF is both necessary and sufficient for basal and TNF-mediated NF-κB activation and protects against apoptosis both *in vitro and in vivo*. Thus, loss of HAF within liver parenchymal cells increases apoptosis, driving progression to HCC in mice without any further dietary or other manipulation, implicating HAF as a tumor suppressor for HCC within pre-neoplastic hepatocytes. In established HCC, HAF levels are increased in malignant hepatocytes and in immune infiltrating cells (**Fig. 6F**), suggesting potential distinct roles of HAF in pre-neoplastic tissue versus HCC.

The effects of genetic HAF deletion in mice closely resemble that of deletion of key components of the NF-κB pathway. Homozygous deletion of HAF in liver parenchymal cells (Alfp-Cre; LPC-S^-/-^) is embryonic lethal and is reminiscent of that seen with homozygous deletion of p65 (RelA) in mice which is also embryonic lethal due to massive TNF-induced hepatocyte apoptosis (56-58). Homozygous deletion of HAF in hepatocytes (Alb-Cre; hepS^-/-^) was tolerated presumably due to the incomplete deletion of HAF (**Fig. 1B, 3C**) but resulted in tumors with hallmarks of MASH in 42% of male and 25% of female mice at age 18 months. Knockout of NEMO in liver parenchymal cells similarly resulted in MASH-like features alongside increased apoptosis ultimately resulting in HCC (59). Additionally, mice with double knockout of RIPK1 and TRAF2 showed impaired NF-κB activation, increased apoptosis followed by HCC development (60). These studies suggest that deficiency in HAF and/or NF-kB activation promotes hepatocarcinogenesis at least in part, by enhancing pro-inflammatory cell death. Paradoxically, constitutive activation of the NF-κB pathway is commonly observed in HCC and in later stages of tumor development and NF-κB inhibition at this stage does not impact tumor initiation but instead prevents progression to HCC (18, 65). Similarly, we show that HAF levels are suppressed in livers with simple steatosis but increased during steatohepatitis/MASH/cirrhosis/HCC from both mice and humans which, in mice, mirror P-p65 and TRADD levels (**Fig. 6E-F**). This suggest that HAF, similar to NF-κB, can act either as as tumor suppressor or tumor promote depending on cellular context.

Although previous studies have implicated HAF in the regulation of the NF-κB pathway, our study presented here reveals for the first time, a mechanism for the regulation of NF-κB signaling by HAF (66, 67). The transcriptional regulation of TRADD and RIPK1 identified in this study adds to a growing list of validated HAF transcriptional targets (*EPO, VEGFA, POU5F1*), which complement HAF’s other functions including as an atypical E3 ubiquitin ligase for HIF-1α and neurofibromin, and as an essential component of the spliceosome (62). Due to its multifunctional properties, it is likely that HAF may mediate its effects on the NF-κB pathway at multiple levels. Related to this, the spliceosome interacting partner of HAF, RSRP1, was shown to promote the mesenchymal phenotype in glioblastoma cells through NF-κB activation mediated through alternative splicing (68). Thus, it is possible that HAF deficiency *in vitro* and *in vivo* may also result in splicing and other defects independent of the NF-κB pathway that contribute to apoptosis and turnover that leads to HCC. Indeed, we show that HAF knockdown promotes the alternative splicing of the anti-apoptotic Bcl-X_L_ protein to favor its pro-apoptotic, Bcl-X_S_ form (**Fig. 5C, S5G**), which may contribute to the increased apoptosis within HAF-deficient cells independent of the NF-κB. Related to this, RIPK1 and TRADD have also been shown to undergo alternative splicing, which decreases protein abundance and may give the appearance of transcriptional downregulation depending on the primer/probe sets employed for qPCR analysis (55). However, our analysis using exon-specific primers did not detect the presence of alternative exons in *TRADD* and *RIPK1* (**S4F-G**) suggesting that the decrease in TRADD and RIPK1 observed with HAF deficiency was primarily associated with decreased transcription, rather than defective splicing. Regarding HAF’s other role as an E3 ubiquitin ligase for HIF-1α, we did not observe any induction of HIF-1α within pre-neoplastic hepatocytes of hepS^-/-^ or LCP-S^+/-^ mice independent of hypoxia (**Fig. 6B**) suggesting that HIF-1α upregulation is unlikely to be involved in the hepatocarcinogenesis associated with HAF loss.

In summary, our findings reveal a novel role of HAF in preventing hepatocyte apoptosis, thus protecting against progression to HCC in the context of MASH in part through the transcriptional regulation of the NF-κB pathway. HAF levels are suppressed in livers with simple steatosis but increased during HCC in humans suggesting a context-specific role of HAF and its potential utility as a biomarker for progression from simple steatosis to MASH and HCC. Additionally, the targeting of HAF in advanced MASH/established HCC could be a strategy to fine-tune NF-κB activity (and of HAF’s other pathological targets) to preserve NF-κB’s physiological function while restraining its pathological role.

## Acknowledgements and funding sources

Supported by R21DK115991-NIH-NIDDK, R01DK128819-NIH-NIDDK and UL1TR002538-NIH-NCATS to PNM; R35GM150766-01-NIH-NIGMS to MG; R01CA222570-NIH-NCI, Damon Runyon-Rachleff Innovation Award DR 61-20 and ACS grant RSG-22-014-01-CCB to KJE; and R01CA262262-NIH-NCI to MYK. This work was supported by UROP from the Office of Undergraduate Research at the University of Utah awarded to LM-TH. We acknowledge financial support by the Huntsman Cancer Foundation and the University of Utah Mutation Generation and Detection Core and the Transgenic and Gene Targeting Mouse Core for generating the SART1 conditional mouse line used in this study (funded by U54DK110858) and the Metabolomics Core at the University of Utah. The content is solely the responsibility of the authors and does not necessarily represent the official views of the NIH.

## Notes

**Conflict of interest:** MYK is founder and has equity in Kuda Therapeutics Inc. All other authors declare no conflict of interest.

### Competing Interest Statement

MYK is founder and has equity in Kuda Therapeutics Inc. All other authors declare no conflict of interest.

