## Supplemental Figs and Legends for "HAF Prevents Hepatocyte Apoptosis and Hepatocellular Carcinoma through Transcriptional Regulation of the NF-κB pathway"

**Supplemental figure legends:**

**S1:** A) Quantitative RT-PCR analysis of *SART1* mRNA expression levels of control and HepS$^{-/-}$ mice (n=6 mice per group. B) Liver transaminase, ALT levels in male and female control (n=4 males, n=3 females) and HepS$^{-/-}$ (n=4 males, n=3 females) mice at 6 months of age. C) NASH-activity scores (NAS) of female control and HepS$^{-/-}$ mice at 18 months. Data represents mean ± SEM. Each data point indicates score from an individual mouse. Red shapes indicate tumors. D) Tumor incidence in female hepS$^{-/-}$ versus their wild type litter-mates (Con) at 18 months. E) Gross images of livers from hepS$^{-/-}$ female mice showing tumors. Scale bar: 1 cm. F) Representative H&E of livers from 18-month-old female Con and hepS$^{-/-}$ mice showing HCC in hepS$^{-/-}$ liver.

**S2:** A) Levels of most abundant acylcarnitines (ACars) in livers of 6-month-old male control (grey bars) and hepS$^{-/-}$ (blue bars) mice (5 mice/group). B) Levels of d18:1 ceramides in livers or liver tumors from 18-month-old male control and hepS$^{-/-}$ mice (5 mice per/group except tumors which are 3 mice per group). Data are the mean ± SEM.

**S3**: A) Western blot quantification of HAF and NF-kB pathway components in pooled isolated primary hepatocytes from control (n=2) and LPC-S$^{+/-}$ mice (n=2) after TNF treatment. B) Western blot showing effects of siRNA HAF knockdown on the NF-kB pathway in Snu475 cells. C) Immunofluorescence of HAF and P-p65 in Snu475 cells transfected with HAF siRNA. D) Western blot showing HepG2 cells transfected with HAF siRNA for three different time points and its impact on the NF-kB pathway.

**S4**: A) Western blot of HAF and NF-kB pathway components in isolated hepatocytes pooled from two 3-month-old control and two LPC-S$^{+/-}$ mice respectively. B) Western blot showing protein levels of TRADD, RIPK1, TAK1 and NEMO after indicated durations of treatment with 25 ug/ml of cycloheximide added 24hrs after HAF siRNA transfection in HepG2 cells. Western blot is representative of three independent experiments. C) Western blot showing effects of treatment of HepG2 cells with 30uM lysosome inhibitor chloroquine (4hrs) added 48hrs after siRNA transfection. D) Quantitative RT-PCR of NEMO and TAK1 mRNA levels in HEPG2 cells transfected with HAF siRNA. E) Luciferase reporter activity of the TRADD site 1 (TS1) reporter construct compared to that of a similar reporter construct bearing a deletion of the HAF binding site within TS1 (TD1) in ACHN cells. F) qRT-PCR to detect alternative splicing of TRADD and RIPK1 in HepG2 cells transfected with HAF siRNA with alternatively spliced isoforms/ variants shown inset.

**S5:** A) Results from resazurin cell viability assay in THLE3 cells transfected with two independent HAF siRNAs. B) Annexin V staining in HUH7 cells transfected with HAF siRNA. C) Western blot showing time course effects of HAF siRNA transfection in HEPG2 cells on cleavage of caspase 3. D) Cell viability assessed by resazurin assay in HEPG2 cells transfected with HAF siRNA for 72hrs then treated with TNF for 24hr. E) Semi-quantitative PCR showing alternative splicing of Bcl-$X_L$/Bcl-$X_S$ in Huh7 cells transfected with two independent HAF siRNAs. F) Annexin V staining in HEPG2 cells transfected with HAF siRNA for 48hrs then treated with TNF for 8hr. G) Western blot of livers showing alternative splicing of Bcl-$X_L$/Bcl-$X_S$ in whole livers harvested from 6-month-old male control and LPC-S$^{+/-}$ mice with densitometric quantitation shown in H. I) Annexin V staining of isolated hepatocytes from 6-month-old control and LPC-S $^{+/-}$ mice. Staining reagent was added 3hrs after hepatocyte isolation and read 6hrs later.

**S6:** Western blots of livers of C57BL/6 mice fed a chow diet for 4, 24, 48 and 72 weeks.

### Supplementary Data S1

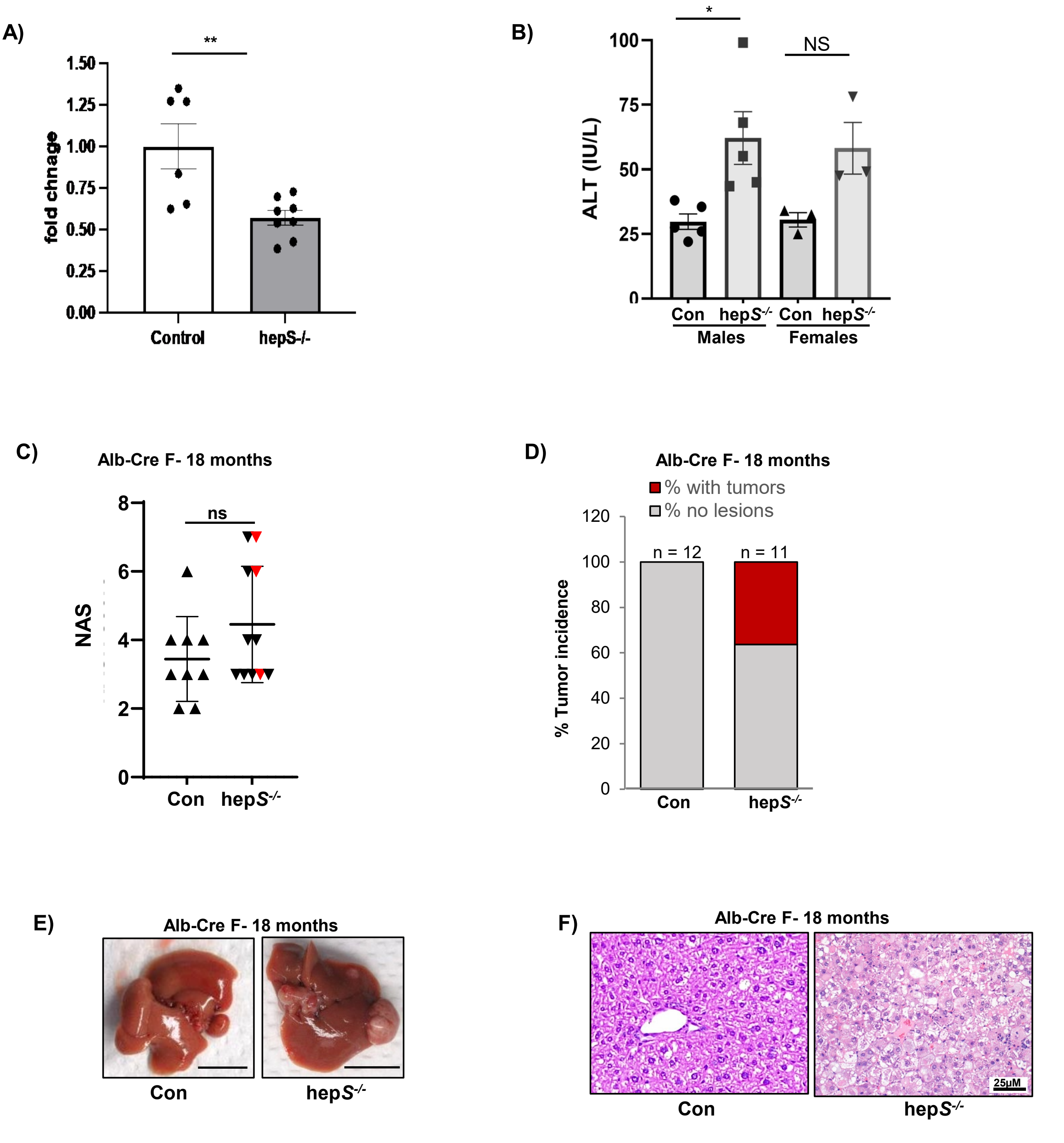

### Supplementary Data S2

A)

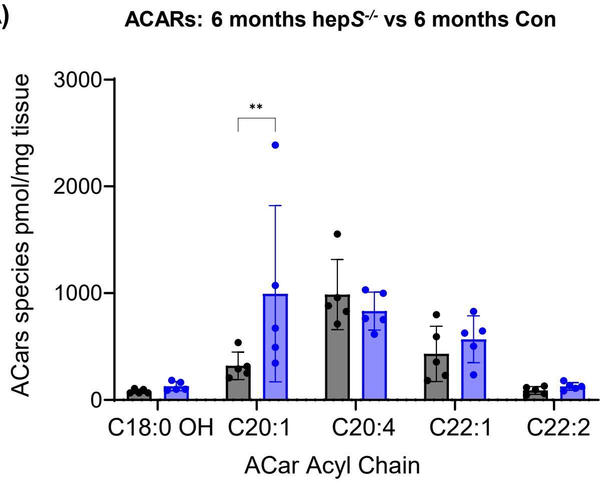


B)

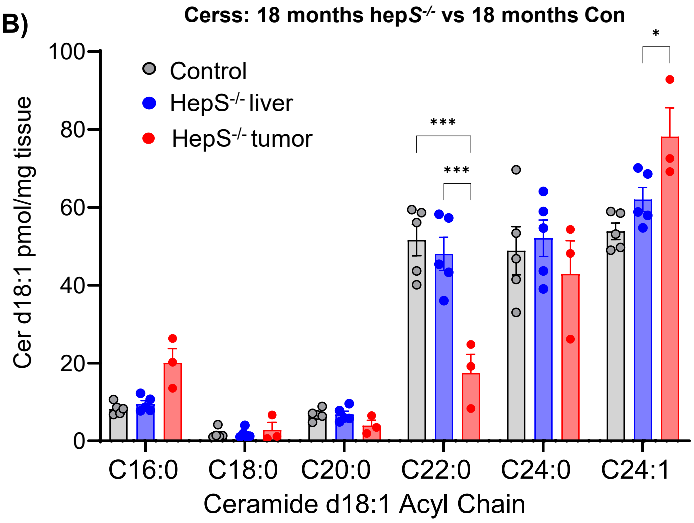

### Supplementary Data S3

A)

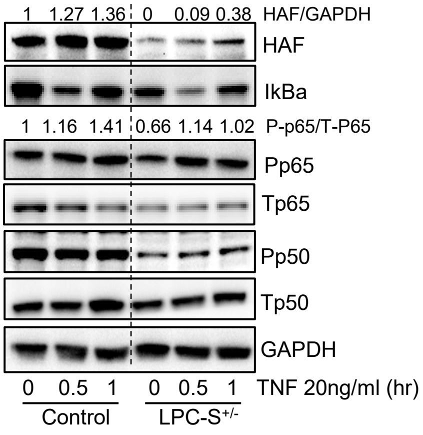


B)

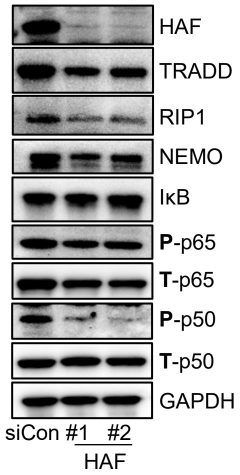


C)

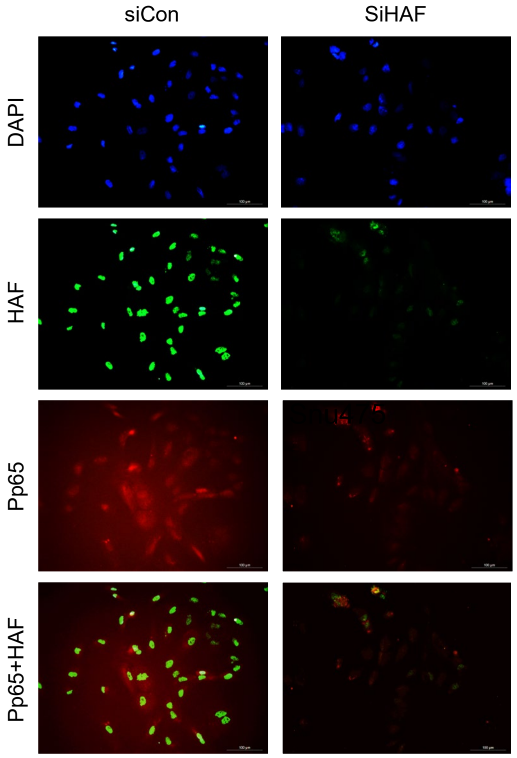


D)

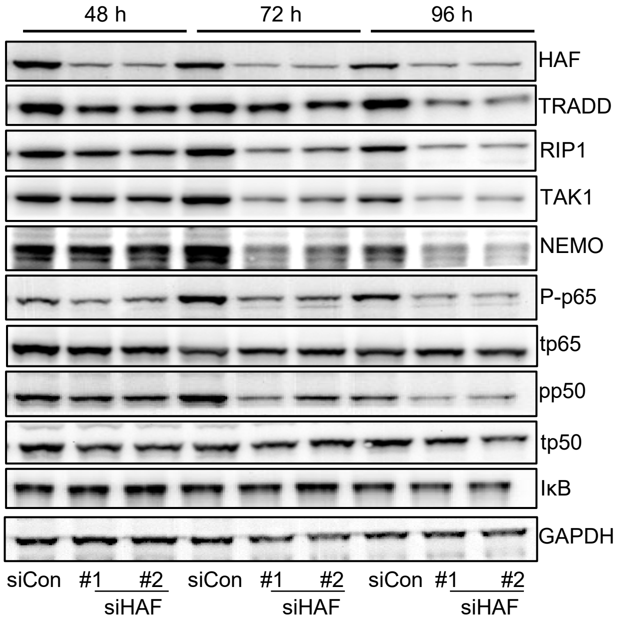

### Supplementary Data S4

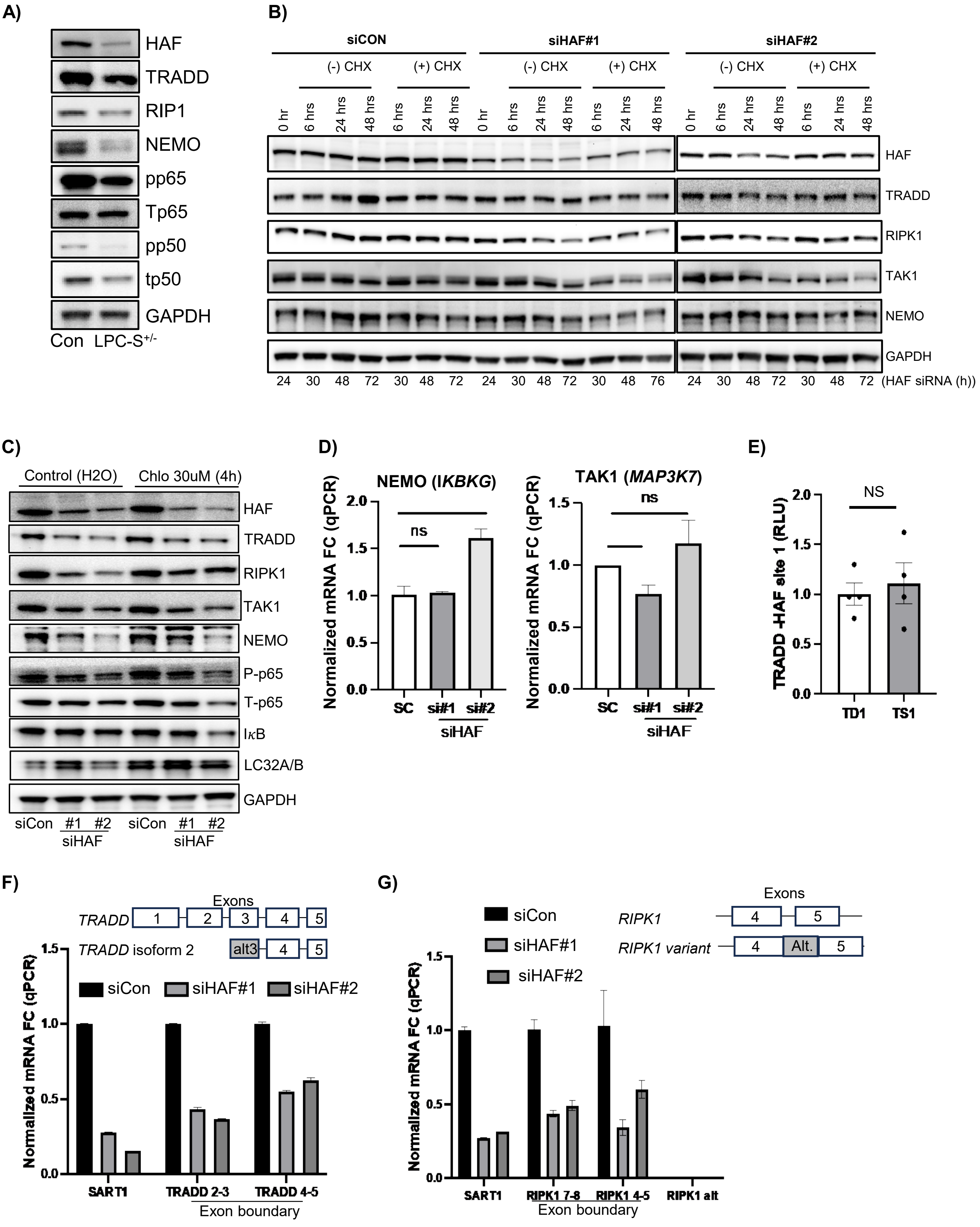

### Supplementary Data S5

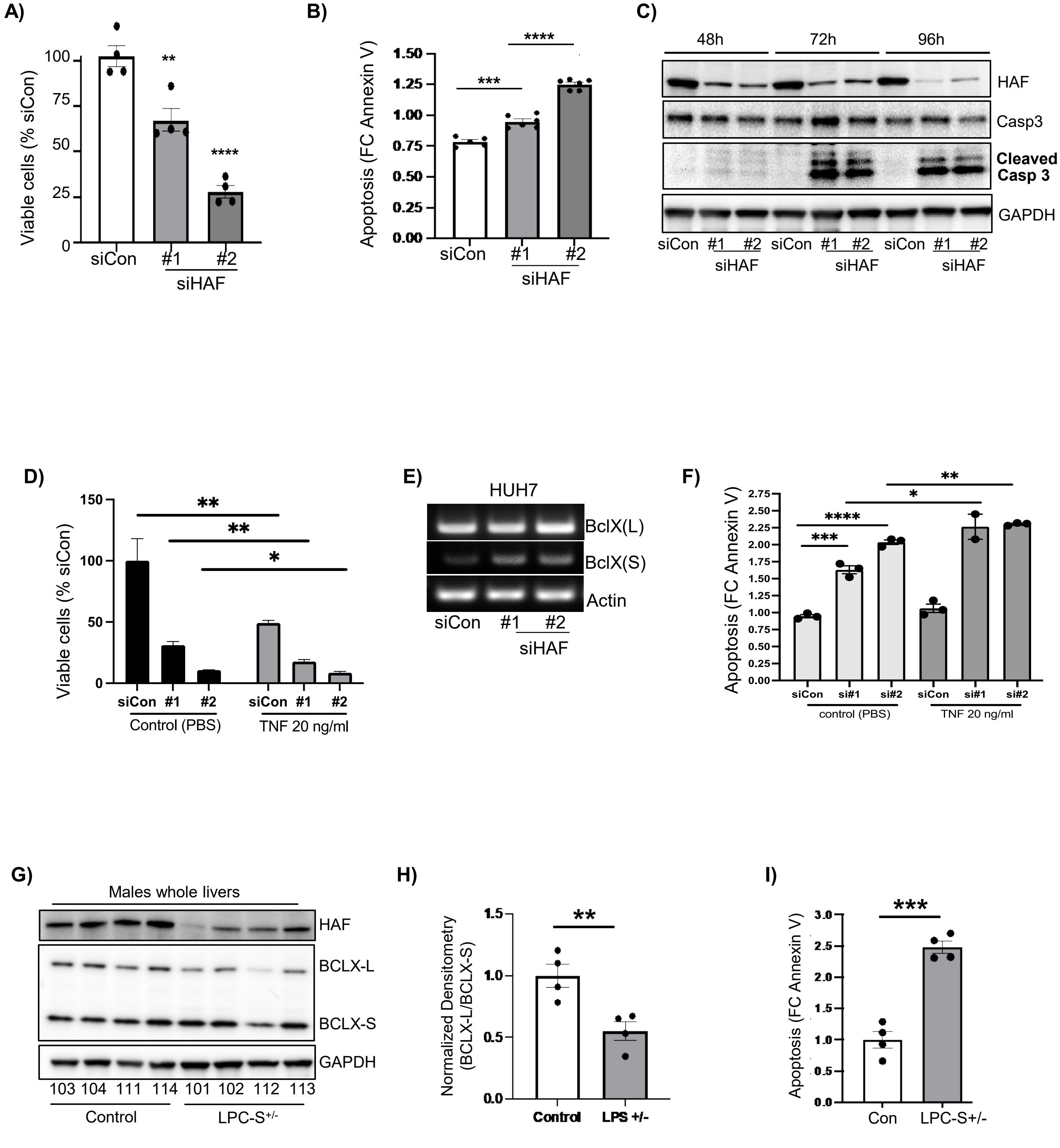

**A)**

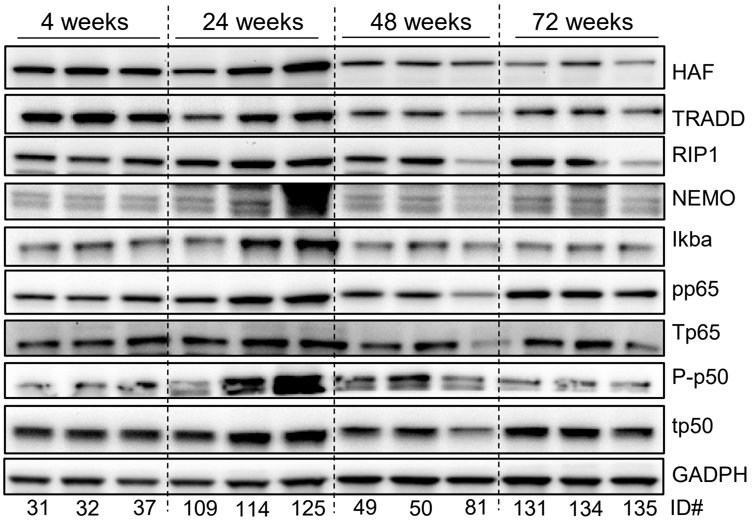
4 weeks
24 weeks
48 weeks
72 weeks
HAF
TRADD
RIP1
NEMO
Ikba
pp65
Tp65
P-p50
tp50
GADPH
31
32
37
109
114
125
49
50
81
131
134
135
ID#

#### Method Set: D:\Data\Golkowski_Lab\Thankhoe \20230915_TR_LS_HEK_1_DDA_Slot1-30_1_1942.d\1942.m

### otofControl

#### General Information

**Method Name:** DDA_Long_Gradien_45min.m
**Application Name:** timsTOF engine
**Device Type:** timsTOF Pro 2
**Operator:** unknown
**Operating System:** unknown

**Saved:** 2023/08/03 15:29:49/07:00
**Application Version:** 4.1.4.12
**Device Serial Number:** 1875087.10665
**Host:** unknown
**Organisation:** Bruker Daltonik GmbH

#### Chromatogram TIMS

##### Chromatogram Traces TIMS

| Enabled | Color | Type | Masses | Width | Polarity | Filter | Mobility |
|---|---|---|---|---|---|---|---|
| On | Dark Blue | BPC | | | ± | MS | |
| On | Red | TIC | | | ± | All MS/MS | |
| On | Green | TIC | | | ± | MS | |

#### Mobilogram

##### Mobilogram list

| Enabled | Color | Type | Masses | Width | Polarity | Filter |
|---|---|---|---|---|---|---|
| On | Blue | EIM | 922 | ±0.05 | + | ALL |
| On | Green | EIM | 1222 | ±0.05 | + | ALL |
| On | Red | EIM | 622 | ±0.05 | + | ALL |

#### Calibration TOF

**Ion Polarity:** Positive
**Calibration Date:** 2023/09/14 14:56:10/07:00
**Calibration Mode:** Linear
**Ion Polarity:** Negative
**Calibration Date:** 2022/07/14 14:21:24+01:00
**Calibration Mode:** Enhanced Quadratic

**Operator:** unknown
**Ref. Mass List:** Tuning TOF
**Score:** 100.00 %
**Operator:** Demo User
**Ref. Mass List:** Tuning Mix ES-TOF (ESI)
**Score:** 100.00 %

#### Calibration TIMS

**Ion Polarity:** Positive
**Calibration Date:** 2023/09/14 15:00:12/07:00
**Calibration Mode:** Linear
**Ion Polarity:** Negative
**Calibration Date:** 2022-07-01T10:12:13+01:00
**Calibration Mode:** Linear

**Operator:** unknown
**Ref. Mass List:** Tuning TIMS
**Score:** 100.00 %
**Operator:** Demo User
**Ref. Mass List:** Tuning Mix ES-TOF (ESI)
**Score:** 100.00 %

#### SPL

**Scheduled Precursor List:** Off

#### Segment 1

#### 0 .... 5 min

#### Main

| | | | |
|---|---|---|---|
| **Polarity:** | Positive | **TIMS enable:** | On |
| | | **Scan Mode:** | PASEF |
| **Mass Range from:** | 100 m/z | **Mass Range to:** | 1700 m/z |
| **1/K0 Start:** | 0.60 Vs/cm² | **1/K0 End:** | 1.40 Vs/cm² |
| **Rolling Average:** | On | **Rolling Average No.:** | 10 |
| **View:** | Expert | | |

#### Mode General TIMS

| | | | |
|---|---|---|---|
| **Enable TIMS:** | On | **Mark as Calibration Segment:** | Off |
| **MS Data Reduction:** | Off | **Save Spectra:** | Save Frame |
| **Mass Spectra Peak Detection:** | Use Max. Intensity | **Absolute Threshold:** | 10 |
| **Intensity Threshold:** | Absolute | **Intensity Threshold:** | 5000.00 |

#### Mode TIMS

| | | | |
|---|---|---|---|
| **ICC:** | Off | **Target:** | 2000000.0 Mio. |
| **imeX mode:** | Custom | **Resolution:** | Custom |
| **1/K0 Start:** | 0.60 Vs/cm² | **1/K0 End:** | 1.40 Vs/cm² |
| **Ramp Time:** | 120.0 ms | **Spectra Rate:** | n/a Hz |
| **Advanced Parameters:** | On | | |
| **Lock accumul. to mob. range:** | On | **Lock Duty Cycle to 100 %:** | On |
| **Accumulation Time:** | 2.0 ms | **Duty Cycle:** | n/a % |
| **Cycle Time:** | n/a ms | | |

#### Source

| | | | |
|---|---|---|---|
| **Source:** | CaptiveSpray | | |
| **Capillary:** | 1600 V | **NanoBooster:** | Off |
| **Dry Gas:** | 3.0 l/min | **Dry Temp:** | 180 °C |
| **Divert Valve:** | Waste 1-6 | | |

#### Tune General

| | | | |
|---|---|---|---|
| **Funnel 1 RF:** | 300.0 Vpp | **isCID Energy:** | 0.0 eV |
| **Deflection Delta:** | 70.0 V | **Funnel 2 RF:** | 200.0 Vpp |
| **Multipole RF:** | 500.0 Vpp | **Ion Energy:** | 5.0 eV |
| **Low Mass:** | 200.00 m/z | **Collision Energy:** | 10.0 eV |
| **Collision RF:** | 1500.0 Vpp | **Transfer Time:** | 60.0 µs |
| **Pre Pulse Storage:** | 12.0 µs | **Stepping:** | Off |

#### Tune TIMS

| | | | |
|---|---|---|---|
| **D1 (DeflTransf -> CapEx):** | -20.0 V | **D2 (DeflDisc -> DeflTransf):** | -160.0 V |
| **D3 (Fun1In -> DeflTransf):** | 110.0 V | **D4 (AccuTrap -> Fun1In):** | 110.0 V |
| **D5 (AccuEx -> AccuTransf):** | 0.0 V | **D6 (RampStart -> AccuExit):** | 55.0 V |
| **Funnel1 RF:** | 450.0 Vpp | **Collision Cell In:** | 300.0 V |

## MS/MS

##### *PASEF*

| | | | |
|---|---|---|---|
| **PASEF:** | On | **No. of PASEF MS/MS scans:** | 10 |
| **Charge Range Min.:** | 0 | **Charge Range Max.:** | 5 |
| **Active Exclusion:** | On | **Release after:** | 0.40 min |
| **Reconsider Precursor,if:** | On | **Curr. Intens./Prev.Intens.:** | 4.00 |
| **Target Intensity:** | 5000 cts/s | **Intensity Threshold:** | 1000 cts/s |
| **Use Intens./Repet.Table:** | Off | | |

##### *PASEF CID*

| | | | |
|---|---|---|---|
| **Advanced Coll.Energy Settings:** | Off | | |
| **Isol.Mass Start:** | 700.00 m/z | **Isol.Mass End:** | 800.00 m/z |
| **Isol.Width Start:** | 2.00 m/z | **Isol.Width End:** | 3.00 m/z |
| **1/k0 Start:** | 0.60 V s/cm² | **1/k0 End:** | 1.60 V s/cm² |
| **Energy Start:** | 20.00 eV | **Energy End:** | 59.00 eV |

| | |
|---|---|
| **TIMS Stepping:** | Off |

##### PASEF Advanced

| | | | |
|---|---|---|---|
| **Pasef MS Repetitions:** | 1 x | **Overlap:** | 4 |
| **Summation Width:** | 25 pts. | **Max. No. of Peaks:** | 3 |
| **Mass Width:** | 0.015 m/z | **1/K0 Width:** | 0.015 V s/cm² |

##### Scheduling List

| Intensity | Repetition |
|---|---|
| 0 | 10 |
| 6324 | 10 |
| 6666 | 9 |
| 7071 | 8 |
| 7559 | 7 |
| 8164 | 6 |
| 8944 | 5 |
| 10000 | 4 |
| 11547 | 3 |
| 14142 | 2 |
| 20000 | 1 |

##### *MRM*

| | |
|---|---|
| **MRM:** | Off |

##### *bbCID*

| | |
|---|---|
| **bbCID (MS-MS/MS):** | Off |

##### Processing

| | |
|---|---|
| **Processing Classic:** | Off |

#### Processing TIMS

| | | | |
|---|---|---|---|
| **Processing TIMS:** | On | | |
| **Absolute Threshold:** | 10.0 cts. | **Abs.Thr.(per 100 sum. Accu Time):** | n/a |
| **Use Maximum Intensity:** | On | **Sensitivity:** | 99.00 % |
| **Area Threshold:** | Off | **Area Threshold:** | -- |
| **Intensity Threshold:** | Absolute | **Intensity Threshold:** | 5000.00 |
| **Min. Peak Valley:** | On | **Min. Peak Valley:** | 10.00 % |
