## Supplemental methods for "HAF Prevents Hepatocyte Apoptosis and Hepatocellular Carcinoma through Transcriptional Regulation of the NF-κB pathway"

*siRNA and other reagents.* The siRNA used were ON-Target plus SMART pool (DharmaconL-017283, siHAF#1), HS Sart1 4 flexiTube (Qiagen S100710808, siHAF#2), RIP1 SMARTpool (Dharmacon M-004445-02) and TRADD SMARTpool (Dharmacon M-004452-01) at 10nM final concentrations. DNA plasmids used were pCMV14-3xFLAG-HAF was as previously described (1), whereas siRNA resistant pCMV14-3xFLAG-HAF(r) was generated based by synonymous mutations of the siHAF#2 binding sequence (binding sequence: CAGCATCGAGGAGACTAACAA, resistant sequence: CAGTATAGAAGAAACCAATAA). Other constructs were purchased: TRADD (HG13888-NY, Sino Biological Inc, Wayne PA), RIPK1 (HG11268-NY, Sino Biological Inc) and NEMO (Addgene 13512, Watertown, MA). The TRADD and RIPK1 SART1 sites (HBS + flanking sequences) and SART1 del (flanking sequences only) were generated by cloning 3 tandem repeats of indicated sites upstream of luciferase and a minimal CMV promoter (pGL4.17, Promega Corp, Madison WI).

*Immunofluorescence.* Snu475 cells were seeded in 8-well chamber slides (Lab-TEK II System 154534, ThermoFisher Scientific), and fixed using 4% formaldehyde for 10 min at room temperature (RT). Cells were washed using Dulbecco’s phosphate buffered saline (DPBS Ca+Mg+, ThermoFisher Scientific). Cell lysis was performed for 10 min at RT with 0.5% Triton diluted in phosphate buffered saline (PBS Ca-Mg-, ThermoFisher Scientific), followed by blocking using a solution containing DPBS, 3% BSA and 1% normal goat serum (NGS) for 1 h. Incubation with HAF (mouse) and P-p65 antibodies (Cell Signaling Technology) was done overnight at 4C, while the secondary antibodies Alexa-Fluor^TM^ 488 and Alexa-Fluor^TM^ 568 (Invitrogen) were added for 1h at RT. Images were taken with the Leica DM4000 B fluorescent microscope (Leica Microsystems, Deerfield, IL), and processed using Image J.

*Luciferase assay.* NF-kB Luciferase Reporter HepG2 stable cell line and HAF overexpressing ACHN cells were seeded in 96-well plates. For HAF siRNA knockdown and HAF overexpression experiments cells were incubated for 96 h or 48 h, respectively. In some cases, cells were treated with 20 ng/ml of TNF for 4 h prior to the assessment of the relative luciferase activity. The latter was measured using the dual-glo luciferase assay system (Promega, Madison WI) following the manufacturer’s instructions.

*Quantitative Reverse Transcription Polymerase Chain Reaction (qRT-PCR).* Total RNA was isolated from HepG2 cells using RNeasy Kit (Qiagen) and quantified by NanoDrop 2000c (ThermoFisher Scientific). RNA samples were prepared for reverse transcription using 1ug of RNA, RT enzyme and buffer mix. qRT-PCR reactions were performed using TaqMan^TM^ Universal PCR master mix (Applied Biosystems) and the following probes SART1 (HS00193002_m1, Mm00600280), TRADD (Hs00182558_m1, Hs00601065_g1), RIPK1 (Hs01041869_m1, Hs01041868_m1, APTZ9NC), NEMO (HS00415849_m1), TAK1 (Hs00177373_m1) and B2M (Hs00187842_m1, Mm00437752) from ThermoFisher Scientific. Reactions were run on Light Cycler® 480 (Roche). Expression levels were determined based on ΔΔ Ct calculations relative to siControl or empty vector.

*Generation of SART1 floxed mice using EASI-Crispr.*

The Easi-CRISPR (Efficient additions with ssDNA inserts-CRISPR) method, which is a well-established and highly-efficient method to mediated targeted genomic knockins, was used to generate a floxed Sart1 allele (2). Mouse embryos were injected with two preassembled guide RNA and Cas9 endonuclease ribonucleoprotein (RNP) complexes to create double stranded DNA breaks around exons 4 and 5. A long single-stranded DNA (lssDNA) donor engineered to contain a floxed version of exons 4 and 5 (grey triangles), flanked by 100- base pair left and right homology arms (blue) was co-injected to serve as a DNA repair template. The guide RNAs were designed to direct Cas9 mediated cutting immediately adjacent to each homology arm, which facilitated insertion of the lssDNA donor via homology directed repair (HDR). This results in replacement of the wildtype exons 4 and 5 with a floxed version. Insertion of the floxed allele was identified via simple PCR and restriction fragment length polymorphism (RFLP) analyses.

*Isolation of mouse peritoneal macrophages*. Peritoneal macrophages were isolated from Control or macS^-/-^ mice as described (3). Briefly, mice were injected with 2.5ml of sterilized 3%(w/v) thioglycollate broth (Difco) within the peritoneal cavity. Macrophages were isolated from the peritoneal cavity 3 days after instillation of thioglycolate as described. Isolated cells displayed a viability of >98% as demonstrated by trypan blue exclusion.

**Primer Sequences**

| **Primer Name** | **Sequence 5’ – 3’** | **Utility** |
| --- | --- | --- |
| 3F | agt tgc cag ggt ggt ctg tg | 3 primers used for mouse SART1 locus. Amplicons: 449bp (recombined), 335bp (conditional), 295bp (wild-type) |
| 3R | gcc act caa ctc tgg gat gc |  |
| 5F | tca gtg gga atc tgg gag gt |  |
| 20239 | TGC AAA CAT CAC ATG CAC AC | Alb-Cre: WT (No-Cre) |
| 20240 | TTG GCC CCT TAC CAT AAC TG | Alb-Cre: Common primer |
| oiMR5374 | GAA GCA GAA GCT TAG GAA GAT GG | Alb-Cre: Mutant (Alb-Cre positive) |
| oIMR3066 | CCC AGA AAT GCC AGA TTA CG | LysM-Cre: Mutant (LysM-Cre positive) |
| oIMR3067 | CTT GGG CTG CCA GAA TTT CTC | LysM-Cre: Common Primer |
| oIMR3068 | TTA CAG TCG GCC AGG CTG AC | Lys-Me Cre: WT (no-Cre) |
| Alf-pCre 001 | GTC CAA TTT ACT GAC CGT ACA | Generic primer for Cre detection (sense) |
| Alf-pCre 002 | CTG TCA CTT GGT CGT GGC AGC | Generic primer for Cre detection (antisense) |

**Lipidomics analysis**

*Chemicals****.*** LC-MS-grade solvents and mobile phase modifiers were obtained from Honeywell Burdick & Jackson, Morristown, NJ (acetonitrile, isopropanol, formic acid), Fisher Scientific, Waltham, MA (methyl *tert*-butyl ether) and Sigma–Aldrich/Fluka, St. Louis, MO (ammonium formate, ammonium acetate). Lipid standards were obtained from Avanti Polar Lipids, Alabaster, AL (EquiSPLASH LIPIDOMIX (330731)) and Cayman Chemical, Ann Arbor, MI, (Palmitic acid-d31 **#**16497).

*Sample Preparation.* Extraction of lipids was carried out using a biphasic solvent system of cold methanol, methyl *tert*-butyl ether (MTBE), PBS, and water as described with some modifications (4). In a randomized sequence, to each sample was added 225 µL MeOH with internal standards and 188 µL PBS. Samples were homogenized for 30 seconds, transferred to microcentrifuge tubes (polypropylene 1.7 mL, VWR, USA) containing 750 µL MTBE, and then incubated on ice with occasional vortexing for 1 hr. Following incubation, samples were centrifuged at 15,000 x g for 10 minutes at 4 °C. The organic (upper) layer was collected, and the aqueous (lower) layer was re-extracted with 1 mL of 10:3:2.5 (*v/v/v*) MTBE/MeOH/dd-H2O, briefly vortexed, incubated at RT, and centrifuged at 15,000 x g for 10 minutes at 4 °C. Upper phases were combined and evaporated to dryness under speedvac. Lipid extracts were reconstituted in 600 µL of 4:1:1 (v/v/v) IPA/ACN/water and transferred to LC-MS vials for analysis. Concurrently, a process blank sample was prepared and pooled quality control (QC) samples were prepared by taking equal volumes from each sample after final resuspension.

*Mass Spectrometry Analysis of Samples.* Lipid extracts were separated on an Acquity UPLC CSH C18 column (2.1 x 100 mm; 1.7 µm) coupled to an Acquity UPLC CSH C18 VanGuard precolumn (5 × 2.1 mm; 1.7 µm) (Waters, Milford, MA) maintained at 65 °C connected to an Agilent HiP 1290 Sampler, Agilent 1290 Infinity pump, and Agilent 6545 Accurate Mass Q-TOF dual AJS-ESI mass spectrometer (Agilent Technologies, Santa Clara, CA). Samples were analyzed in a randomized order in both positive and negative ionization modes in separate experiments acquiring with the scan range m/z 100 – 1700. For positive mode, the source gas temperature was set to 225 °C, with a drying gas flow of 11 L/min, nebulizer pressure of 40 psig, sheath gas temp of 350 °C and sheath gas flow of 11 L/min. VCap voltage is set at 3500 V, nozzle voltage 500V, fragmentor at 110 V, skimmer at 85 V and octopole RF peak at 750 V. For negative mode, the source gas temperature was set to 300 °C, with a drying gas flow of 11 L/min, a nebulizer pressure of 30 psig, sheath gas temp of 350 °C and sheath gas flow 11 L/min. VCap voltage was set at 3500 V, nozzle voltage 75 V, fragmentor at 175 V, skimmer at 75 V and octopole RF peak at 750 V. Mobile phase A consisted of ACN:H_2_O (60:40, *v/v*) in 10 mM ammonium formate and 0.1% formic acid, and mobile phase B consisted of IPA:ACN:H_2_O (90:9:1, *v/v/v*) in 10 mM ammonium formate and 0.1% formic acid. For negative mode analysis the modifiers were changed to 10 mM ammonium acetate. The chromatography gradient for both positive and negative modes started at 15% mobile phase B then increased to 30% B over 2.4 min, it then increased to 48% B from 2.4 – 3.0 min, then increased to 82% B from 3 – 13.2 min, then increased to 99% B from 13.2 – 13.8 min where it’s held until 16.7 min and then returned to the initial conditions and equilibriated for 5 min. Flow was 0.4 mL/min throughout, with injection volumes of 2 µL for positive and 8 µL negative mode. Tandem mass spectrometry was conducted using iterative exclusion, the same LC gradient at collision energies of 20 V and 27.5 V in positive and negative modes, respectively.

*Analysis of Mass Spectrometry Data.* For data processing, Agilent MassHunter (MH) Workstation and software packages MH Qualitative and MH Quantitative were used. The pooled QC (n=8) and process blank (n=4) were injected throughout the sample queue to ensure the reliability of acquired lipidomics data. For lipid annotation, accurate mass and MS/MS matching was used with the Agilent Lipid Annotator library and LipidMatch (5). Results from the positive and negative ionization modes from Lipid Annotator were merged based on the class of lipid identified. Data exported from MH Quantitative was evaluated using Excel where initial lipid targets are parsed based on the following criteria. Only lipids with relative standard deviations (RSD) less than 30% in QC samples are used for data analysis. Additionally, only lipids with background AUC counts in process blanks that are less than 30% of QC are used for data analysis. The parsed excel data tables are normalized based on the ratio to class-specific internal standards, then to tissue mass and sum prior to statistical analysis.

**Proteomics Analysis**

*Preparation of peptide samples for LC-MS/MS analysis.* Peptide samples were prepared as described previously with a few minor modification (6). Briefly, 250 μL of denaturing lysis buffer (8 M urea, 100 mM Tris, pH = 8.5) containing 5 mM tris(2-carboxyethyl)phosphine hydrochloride (TCEP*HCl, Cat# PI20491) and 10 mM 2-chloroacetamide (CAM, Fisher Scientific, Cat# U15-500) was added to aliquots of mouse liver (~50 mg wet weight) and homogenized with a D1000 Hand-held homogenizer (Benchmark Scientific, Cat# D1000) until homogenous (~10-15 s) on wet ice. Samples were centrifuged at 14,000 rcf for 20 min. at 4°C to pellet insoluble debris, and the protein content determined using the Pierce 660 nm protein assay reagent (Thermo Scientific, Cat# 22660), with a Multiskan Skyhigh Microplate Spectrophotometer (Thermo Scientific). To purify proteins, acetone precipitation was performed. Briefly, protein extracts in 8 M urea were diluted with four times the volume of ice-cold acetone (Alfa Aesar, Cat# 22928-K7), 80% aq. trichloroacetic acid (TCA, Spectrum Laboratory Products, Cat# T-161-500ml) was added to a final concentration of 4%, and the mixture vortexed at max speed for 5s. Proteins were allowed to precipitate overnight at -20°C and then pelleted by centrifugation at 2,000 rcf at 4°C for 10 min. The supernatant was aspirated, and the pellets were washed twice with 1 mL ice-cold acetone, dispersing the pellet in a sonicator bath, and pelleting the protein at 2,000 rcf at 4°C for 10 min each time. Pellets were allowed to dry after the final wash for 2-3 min at RT, 250 μL of denaturing lysis buffer (8 M urea, 100 mM Tris, pH = 8.5) was added, and samples were agitated on a Thermal Shake Touch shaker (VWR) at 37°C for 30 min, or until the pellet was dissolved. The protein content of each sample was determined using the Pierce 660nm assay reagent. For digestion, protein concentrations were normalized across samples, samples were diluted 2-fold with 100 mM triethylammonium bicarbonate buffer (TEAB, pH = 8.5, Supelco, Cat#18597-500ML), and adjusted to pH 8-9 with 1 M aq. NaOH. MS-grade Lys-C (1:100 enzyme-to-protein ratio, Fujifilm Wako) was added. The mixture was agitated on the thermal shaker at 37°C for 2 hours, diluted another 2-fold with 100 mM TEAB, and MS-grade trypsin (1:100 enzyme-to-protein ratio, Thermo Scientific, Cat#PI90057) was added. The samples were then agitated on the thermal shaker at 1,400 rpm at 37°C overnight, acidified with formic acid (Fisher Scientific, Cat# A117-50) to 1% of final concentration and cleared by centrifugation for 10 min at 14,000 rcf at room temperature. Peptides were extracted from the supernatant using Oasis HLB 1cc (30 mg) extraction cartridges. Briefly, cartridges are activated by passing through 200 μL of methanol (MeOH, Alfa Aesar, Cat#22909-K7) followed by 200 μL StageTip Buffer B (80% aq. Acetonitrile (ACN, Thermo Scientific, Cat# 325730025) and 0.1% trifluoroacetic acid (TFA)) and equilibrated with 400 μL 0.1% aq. TFA. Peptide solutions were slowly passed through the cartridges using a syringe and cartridges washed with 400 μL 0.1% aq. TFA. Peptides were eluted from the extraction columns using 300 μL Buffer B. 15 μL of the eluate (total peptide sample) was used for global proteome analysis by LC-MS/MS.

*LC-MS/MS analysis and data processing.* Peptide samples were separated on a Bruker nanoElute 2 nano-flow HPLC system using 15 cm C18 PepSep columns (C18, 15 cm x150 uM x 1.5 µM, Bruker, Cat#1893474), with a 45 min LC gradient of 5-35% buffer B in buffer A. LC buffer A was 0.1% aq. formic acid, and buffer B was 0.1% formic acid in acetonitrile (ACN, all solvents were LC-MS grade). Peptides were analyzed using a timsTOF Pro 2 mass spectrometer equipped with a CaptiveSpray nano ESI source (Bruker) in data-dependent acquisition (DDA) mode; details of the MS/MS method can be found in the supplementary information. MS raw files were searched using MaxQuant/Andromeda (7) version 2.4.2.0 and the canonical *Mus musculus* FASTA file (UP000000589_10090) downloaded from the uniport webpage on July 14^th^, 2023. MaxQuant search parameters: TIMS half width = 10, TIMS step = 3, TIMS resolution = 35000, TIMS min MS/MS intensity = 1.5, TIMS remove precursor = enabled, TIMS collapse MS/MS = enabled; variable modifications included Oxidation (M) and Acetyl (Protein N-term); carbamidomethyl (C) was a fixed modification; Label-Free Quantification (LFQ)(8) was enabled with LFQ min. ratio count = 0, normalization count = classic, Fast LFQ = enabled, LFQ min. number of neighbors = 3, and LFQ average number of neighbors = 6; max. missed cleavages was 2, and enzyme was Trypsin/P; protein, peptide and site FDRs were 0.01 and a score minimum of 40 for modified peptides, 0 for unmodified peptides was applied; a delta score minimum of 17 for modified peptides, 0 for unmodified peptides was applied. and max. charge was 7; the initial match tolerance for MS/MS was 20 ppm and 0.5 Da. MaxQuant output files were processed, statistically analyzed, and clustered using the Perseus software package v1.5.6.0 (9). Human gene ontology (GO) terms (GOBP, GOCC and GOMF) were loaded from the ‘mainAnnot.homo_sapiens.txt’ file downloaded on 02.03.2020. Expression columns were log2 transformed. Potential contaminants, reverse hits, and proteins only identified by site were removed. Reproducibility between LC-MS/MS experiments was analyzed by column correlation (Pearson’s r) and replicates with a variation of r > 0.25 compared to the mean r-values of all replicates of the same experiment were considered outliers and excluded from the analyses. Data imputation was performed using a modeled distribution of MS intensity values downshifted by 1.8 and having a width of 0.2. Differential expression was determined using two-tailed, two sample T-tests with Benjamini-Hochberg correction for multiple hypothesis testing (FDR = 0.05).

*GSEA Analysis.* For gene set enrichment analysis (GSEA), we used the ssGSEA2.0 script in R together with the Gene Ontology: Biological Process (GOBP) gene set of the MSigDB database (‘c5.bp.v7.0.symbols’) according to the published protocol with the following minor modifications (10). To rank gene names, we calculated a compound score using the t-test log2 MS intensity ratio multiplied by the -log10 p-value. The parameters used for GSEA were: sample.norm.type = "none", weight = 1, statistic = "area.under.RES", output.score.type = "NES", nperm = 1e3, min.overlap = 10, correl.type = "z.score", par = T, spare.cores = 1, export.signat.gct = T, extended.output = T. To display GSEA results in the heatmap Figure 2F, we calculated an adjusted normalized enrichment score (NES), by multiplying the NES with the -log10 FDR.
